# Reduced parasite burden in feral honeybee colonies

**DOI:** 10.1101/2022.07.18.500457

**Authors:** Patrick L. Kohl, Paul D’Alvise, Benjamin Rutschmann, Sebastian Roth, Felix Remter, Ingolf Steffan-Dewenter, Martin Hasselmann

**Affiliations:** Department of Animal Ecology and Tropical Biology, Biocenter, University of Würzburg, Würzburg, Germany; Department of Livestock Population Genomics, Institute of Animal Science, University of Hohenheim, Stuttgart, Germany; BEEtree-Monitor, Munich, Germany; Katholische Stiftungshochschule München

**Keywords:** Bee viruses, disease management, horizontal transmission, livestock-wildlife interactions, pathogen spillover, swarming, wild honeybees, *Varroa destructor*

## Abstract

Bee parasites are the main threat to apiculture, and since many parasite taxa can spill over from honeybees (*Apis mellifera*) to other bee species, honeybee disease management is important for pollinator conservation in general. It is unknown whether honeybees that escaped from apiaries (i.e., feral colonies) benefit from natural parasite-reducing mechanisms like swarming or suffer from high parasite pressure due to the lack of medical treatment. In the latter case, they could function as parasite reservoirs and pose a risk to the health of managed honeybees (spillback) and wild bees (spillover). We compared the occurrence of 18 microparasites among managed (N=74) and feral (N=64) honeybee colony samples from four regions in Germany using qPCR. We distinguished five colony types representing differences in colony age and management histories, two variables potentially modulating parasite prevalence. Besides strong regional variation in parasite communities, parasite burden was consistently lower in feral than in managed colonies. The overall number of detected parasite taxa per colony was lower, and Trypanosomatidae, chronic bee paralysis virus, and deformed wing viruses A and B were less prevalent and abundant in feral colonies than in managed colonies. Parasite burden was lowest in newly founded feral colonies, intermediate in overwintered feral colonies and managed nucleus colonies, and highest in overwintered managed colonies and hived swarms. Our study confirms the hypothesis that the natural mode of colony reproduction and dispersal by swarming temporally reduces parasite pressure in honeybees. We conclude that feral colonies are unlikely to contribute significantly to the spread of bee diseases.

## Introduction

Disease transfer between managed and wild animals is a potential source of conflict between the livestock sector and nature conservation (Mysterud & Rolandsen, 2019). Populations managed in intensive animal husbandry can be vulnerable to diseases transmitted from wild populations because high animal densities and low genetic variance increase the risk of epidemics (Gortázar et al., 2007). Conversely, diseases of livestock can become a threat to wildlife, e.g., when managed species introduce diseases into non-native areas (Ayala et al., 2020; Costanzi et al., 2021; Schommer & Woolever, 2008). An interesting example of potentially negative livestock-wildlife interactions is that of the disease associations between managed western honeybees (*Apis mellifera*), wild-living honeybees, and other bee species (Fürst et al., 2014; Ravoet et al., 2014). The Western honeybee is native to Africa, Europe, and Western Asia, and is an important pollinator of wild plants both in its native and introduced range (Dick, 2001; Hung et al., 2018). The species is managed by beekeepers globally to pollinate the flowers of crops and to produce honey and other hive products (Crane, 1990; Rollin & Garibaldi, 2019). Where self-sustaining wild populations have gone extinct, as in many regions of Europe and Western Asia (Pirk et al., 2017; Requier et al., 2019), apiculture also serves to protect the honeybee as a species. However, modern rationalized beekeeping can conflict with conservation (Geldmann & González-Varo, 2018; Iwasaki & Hogendoorn, 2022; Panziera et al., 2022). By preventing swarming and maintaining unnaturally large, continuously breeding colonies; by crowding hives in apiaries, and by seasonally moving colonies between bee yards, apiculture promotes the reproduction and spread of bee parasites (Brosi et al., 2017; Loftus et al., 2016; Martínez-López et al., 2022; Nolan & Delaplane, 2017; Peck & Seeley, 2019; Seeley & Smith, 2015). On a global scale, the transport of hives and their products can expose the bees to entirely novel parasites (Goulson & Hughes, 2015). This was famously demonstrated by the worldwide invasions of western honeybee populations by the ectoparasitic mite *Varroa destructor*, whose natural host was the eastern honeybee *A. cerana* (Kraus & Page, 1995; Traynor et al., 2020; Wilfert et al., 2016). Unfortunately, many bee parasites are not restricted to a single species. Since honeybees inevitably share floral resources with other pollinators, their parasites can spill over to, and harm, populations of other bee species (Burnham et al., 2021; Fürst et al., 2014; Piot et al., 2022; Tehel et al., 2022). Therefore, managing honeybee diseases is important for the conservation of bee pollination services in general, both in the contexts of crop production and ecosystem functioning (Bartlett, 2022; Brosi et al., 2017).

Honeybees are social insects that live in large perennial nests, and as such, they are naturally attractive hosts for a range of parasites including arthropods, fungi, bacteria, and viruses (Schmid-Hempel, 1998). However, parasites are not considered to be a limiting factor for wild honeybee populations under natural conditions (Bailey, 1958; Fries et al., 2006; Fries & Camazine, 2001; Ratnieks & Nowakowski, 1989). An important reason seems to be the regulation of parasite populations within colonies as a side-effect of the bees’ natural cycle of colony reproduction and dispersal (DeGrandi-Hoffman et al., 2017; Loftus et al., 2016). Honeybee colonies reproduce via fission when the old queen and approximately 70% of the workers leave as a swarm to build a new nest in another cavity (Seeley, 2010). The swarming bees do not transfer brood to the new nesting site, and it takes around three weeks until a young queen resumes egg-laying in the old nest; hence, brood production is interrupted in both colony parts and the reproduction of parasites infecting the brood or young workers is halted (Loftus et al., 2016; Royce et al., 1991). Furthermore, the rate at which new parasite species enter wild colonies is most probably lower compared to the situation at apiaries, since their nests are typically widely dispersed in the landscape (Lindström et al., 2008; Nolan & Delaplane, 2017; Seeley & Smith, 2015). Besides these ecological factors, wild honeybee populations are more resilient against disease than managed populations because they are more likely to evolve defences against parasites via natural selection (Neumann & Blacquière, 2017; Pirk et al., 2017). A famous example is a population of non-native wild honeybees, inhabiting forests in the species’ introduced range in the north-eastern USA, which was subjected to rapid evolution upon the arrival of *Varroa destructor* in the 1980s and remains viable (Mikheyev et al., 2016; Seeley, 2017). Not only do wild honeybee populations deliver free pollination services (Chang & Hoopingarner, 1991), but they can also benefit the beekeeping sector when genetic adaptations to (novel) stressors, including parasites, are passed on to the managed population (Pirk et al., 2017). Self-sustaining wild honeybee populations exist both in the species’ native range in Africa (Dietemann et al., 2009) and in its introduced range in Australia (Oldroyd et al., 1997), and parts of the Americas (Seeley, 2007; Winston, 1992).

The situation might differ for wild-living honeybee colonies which are recent escapees from apiaries. For example, in Germany, despite active swarm prevention by beekeepers, thousands of honeybee swarms emigrate from bee yards each spring and colonize tree cavities in managed forests. However, their annual survival rate is far below the threshold required to maintain a self-sustaining population (Kohl et al., 2022). We here refer to such honeybees as “feral” (Daniels & Bekoff, 1989), as opposed to “wild”, regardless of whether the native or introduced range of the species is considered. Feral colonies exist wherever apiculture is practised, and they are probably also widespread in Europe (Browne et al., 2020; Dubaić et al., 2021; Kohl et al., 2022; Kohl & Rutschmann, 2018; Oleksa et al., 2013; Rutschmann et al., 2022; Thompson, 2012). On the one hand, feral colonies might have a lower parasite burden than managed colonies due to the natural parasite-reducing effects of swarming and the dispersal of nest sites. On the other hand, with an evolutionary background of artificial selection and a history of care by beekeepers, feral colonies are likely to be stressed by multiple environmental factors and might develop high parasite loads without medical treatment (Thompson et al., 2014).

Understanding the disease ecology of recently feralised honeybee colonies is important because they create a potential management conflict. Honeybees are usually considered livestock animals and beekeepers are obligated to register their hives, regularly apply miticide treatments, and report infectious diseases to prevent epidemics. If feral colonies carry high loads of parasites, however, they might reinfect managed colonies (spillback), undermining the veterinary measures undertaken to combat disease (Frey & Rosenkranz, 2014; Thompson et al., 2014). When highly infected feral colonies disperse into natural areas, they might also function as vectors for parasites that can spill over to non-*Apis* wild bees (Burnham et al., 2021; Fürst et al., 2014; Piot et al., 2022; Tehel et al., 2022). This situation would suggest management options aiming at preventing feralisation or eradicating feral colonies (Taylor et al., 2007). Conversely, where the Western honeybee is a native species, promoting or re-establishing populations of wild-living honeybees is a legitimate conservation goal (Panziera et al., 2022; Requier et al., 2019). It would then be inappropriate to combat feral colonies since they can be the source of future wild populations.

To assess whether feral honeybees might pose a risk to the health of managed and wild bee populations, we compared the occurrence of 18 microparasite taxa in feral and managed honeybee colonies using qPCR (D’Alvise et al., 2019). We collected colony samples from four regions in southern Germany encompassing both rural and urban landscapes, to cover cases from different environments with potentially different parasite communities (Youngsteadt et al., 2015). By taking all samples within a four-week time window in July, we made sure that variation in parasite communities could not be attributable to seasonal variation (D’Alvise et al., 2019; Faurot-Daniels et al., 2020). We chose this point in time because it is epidemiologically relevant: a high frequency of between-colony robbing behaviour due to nectar scarcity in summer (Garbuzov et al., 2020) increases the risks of parasites transmissions, and parasite loads in summer affect subsequent colony winter survival (Ravoet et al., 2013). In addition to the difference between managed and feral, we distinguished five colony types representing different colony age classes and management histories since these factors potentially modulate parasite prevalence. The collected data allowed us to make a nuanced assessment of the effect of environmental differences between managed and feral honeybee colonies on parasite prevalence.

## Material and Methods

### Sample collection

We collected worker bees from managed colonies (N=74, from 73 apiaries) and feral colonies (N=64, from 63 sites) in four regions in southern Germany (Swabian Alb, counties Coburg and Lichtenfels, county Weilheim-Schongau, and the city of Munich; Fig. 1) in July 2020. In the first three study regions, feral colonies were found during a population-monitoring study that was based on systematic censuses of tree cavities made by the black woodpecker, which are the primary nesting sites of honeybee colonies in managed forests in Germany (Kohl et al., 2022; Kohl & Rutschmann, 2018). We sampled from all detected feral colonies whose recent history was known to us (see next paragraph). In the fourth study region (Munich), we were able to select 15 out of approx. 80 feral honeybee nest sites that had previously been mapped by two of us using a combination of private search and citizen reports collected via the BEEtree-Monitor network (S. Roth and F. Remter, unpublished). We selected the 15 nest sites for sampling based on the criteria of accessibility and knowledge of the recent occupation history, and to assure an even distribution over the city and an equal representation of nests in trees and buildings. To obtain locations of managed colonies, we contacted beekeepers via the local beekeeping organizations and asked for permission to sample bees from their hives. This resulted in a sample of managed colonies that roughly matched the sample of feral colonies in terms of size and spatial distribution (Fig. 1). The sampled feral colonies nested in tree cavities or building walls with entrance heights between 2–18 m above the ground. Except for four of the 64 colonies, which had neighbouring colonies in the same tree or building wall, feral colonies were spatially separated from others. The sampled managed colonies were kept in hives and were usually placed next to other colonies in apiaries. The median number of colonies at the apiaries was 6 (range: 1–28 colonies), which well represents the typical apiary size of German beekeepers, who mostly practice beekeeping as a hobby or sideline occupation (Deutscher Imkerbund e.V., 2020).

**Figure 1.**
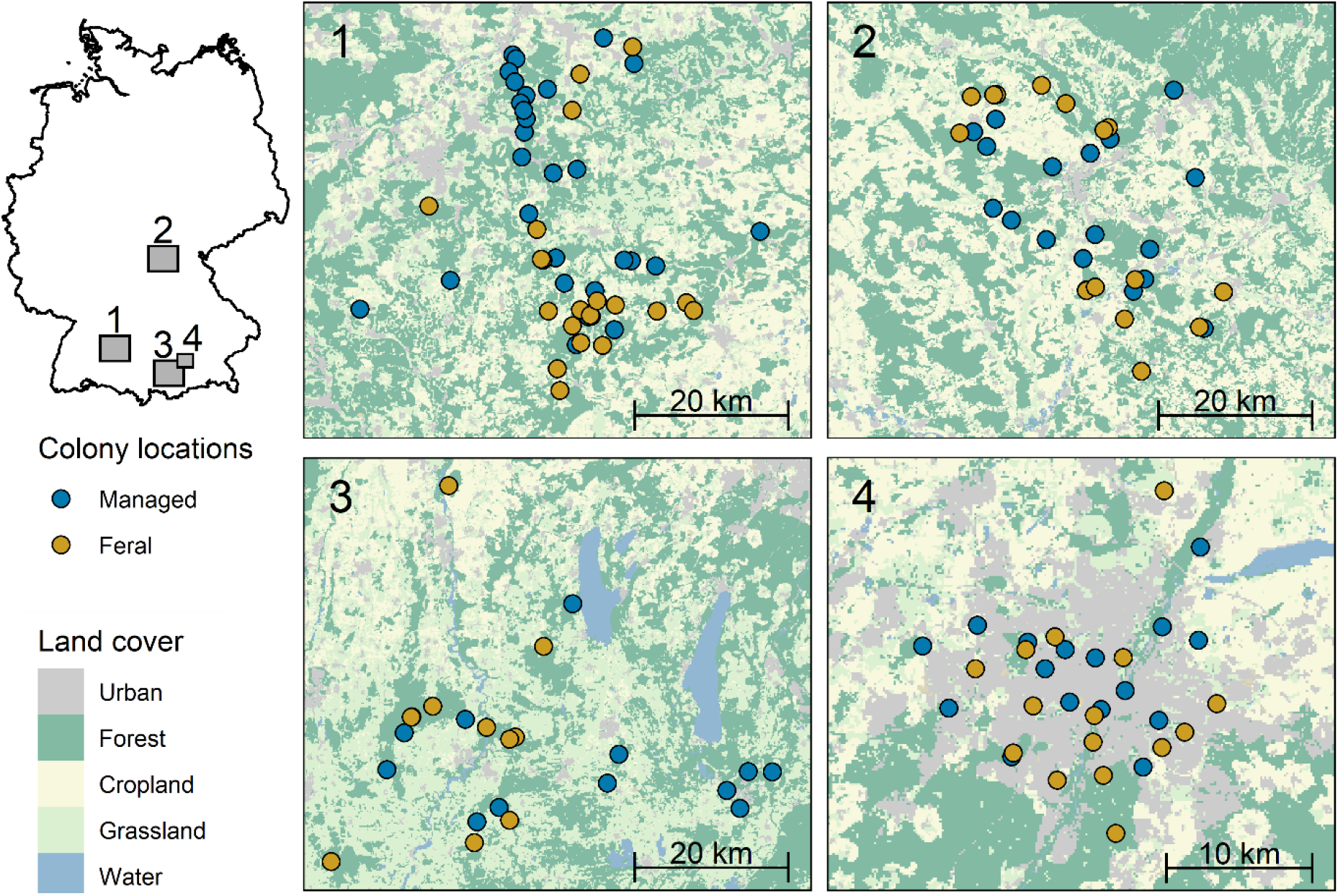
Geographic locations of the sampled managed (N = 74) and feral (N = 64) honeybee colonies in four regions in southern Germany (1: Swabian Alb, 2: Counties Coburg and Lichtenfels, 3: County Weilheim-Schongau, 4: City of Munich). Note that the map of Munich has a different scale. Land cover data are from (Weigand et al., 2020).

A factor potentially affecting parasite burden is colony age. Therefore, we noted for each colony whether it had overwintered at least once (age > 1 year) or had been newly founded in the year of sampling (age < 1 year). Amongst feral colonies 72% (N=46) were recent founders (swarms that moved into the cavity and founded a colony in spring), and 28% (N=18) had overwintered at least once. Amongst managed colonies we sampled similar numbers of overwintered (N=38; 51%) and newly founded colonies (N=36; 49%), since we assumed the ratio of old versus young colonies at apiaries in summer is about 50:50. In the group of young managed colonies, we further distinguished between nucleus colonies, i.e., daughter colonies created by taking brood frames and bees from an established colony (N=27) and hived swarms (N=9). The latter were either natural swarms captured by beekeepers (N=7) or man-made swarms (“packages”) created by brushing bees off the combs of a mother colony and transferring them into a new hive together with a queen (N=2).

Al least 20 bees per colony were captured by placing a white butterfly net in front of the hive or cavity entrance (see supplementary information in (Kohl et al., 2022) for details of the sampling method). With this method, we primarily sampled foragers on their outbound flights. The sampled bees did not show any overt disease symptoms. After capture, they were directly freeze-killed and collected in 50 mL vials. They were permanently kept on dry ice during transport and stored in freezers at −80°C until the analyses.

### Parasite testing

We assessed the colony-level occurrence of 18 microparasites using high-throughput qPCR on a Biomark HD system (Standard BioTools, San Francisco, CA) with parasite-specific primers (Budge et al., 2010; Cepero et al., 2015; D’Alvise et al., 2019; Evans, 2006; Locke et al., 2012; Lourenço et al., 2008; Martínez et al., 2010; Papp et al., 2014) closely following an established protocol (D’Alvise et al., 2019), with the exception that RNA was extracted from homogenates of 20 workers per colony, not from individual bees (see supplementary information for details on RNA extraction and Table S1 for a list of the parasites and reference genes assayed). We ran three qPCR replicates for each assay and considered a parasite taxon as present in a colony if target molecules were detected in at least two. If a colony sample was positive, we averaged the number of target molecules detected in all three assay replicates to obtain a mean number of target molecules per colony sample. As a measure of colony-level parasite abundance, we calculated for each parasite the number of detected target molecules per 100 ng of extracted RNA and reported the logarithm of this ratio, log10(n/100 ng RNA+1), as the “parasite load”.

### Statistical analyses

All statistical analyses were performed with R version 4.0.5 (R Core Team, 2022) and figures were created using “ggplot2” (Wickham, 2016). We analysed whether the number of detected parasite taxa per colony (univariate analysis) and the microparasite community composition (multivariate analysis) differed depending on “management” (two levels: “managed” versus “feral”) and “colony type” (five levels: “managed: overwintered”, “managed: nucleus colonies”, “managed: hived swarms”, “feral: overwintered” and “feral: founders”). In all four analyses, we accounted for the potential effect of spatial location on microparasite communities.

We analysed the number of parasite taxa detected (count data) using generalized linear models (function “glmmTMB”; (Brooks et al., 2017, 2019)). Besides the main factors of interest, “management” or “colony” type”, we included “region” (four levels) as well as the interaction between “region” and “management” or “colony type” as additional predictors to account for spatial differences and for potential variation in the effect of management or colony type between regions. Considering “region” to describe the spatial component was reasonable since most spatial variation in parasite assemblages was attributable to variation between regions rather than variation between sites within regions (see below). Count data can be analysed with a range of model types assuming different probability distributions (Brooks et al., 2017). We therefore first created five models with the same predictor formula but different family functions in “glmmTMB” (family = “poisson”, “nbinom1”, “nbinom2”, “compois” or “genpois”) and selected the best models using the Akaike information criterion for small sample sizes (AICc, function “AICctab” from the “bbmle” package (Bolker, 2022)). In both comparisons (factor “management” or “colony type”) models with a generalized Poisson distribution (family= “genpois”) had the lowest AICc (ΔAICc ≥ 12) so we used these for the analyses. We then tested for significant deviations from model assumptions using the functions “simulateResiduals”, “testResiduals” and “testCategorical” from the “DHARMa” package (Hartig, 2022). No significant deviations were found in tests for uniformity, dispersion, and outliers, but Levene’s test detected significant differences in variance of parasite counts between the four study regions. This problem could be fixed by adding a formula for dispersion with “region” as the single fixed effect. The final formulations of the full models were “glmmTMB(*Number of parasites* ~ X + region + X: region, dispformula = ~ region, family = genpois)” with “X” denoting either “management” or “colony type”. Predictions of the mean number of parasite taxa and 95% confidence limits were produced using the function “emmeans” (Lenth, 2022). To test whether management and colony type or their interactions with region significantly affected parasite counts, we compared pairs of nested models (where one model contained, and the other missed, the predictor of interest) using likelihood ratio tests (LRT, function “anova”). We assessed the effect of management or colony type while accounting for the effect of region (see supplementary information, Tables S2–S4, S11 and S12 for the specifications of the nested models). In the case of the five-level factor “colony type”, we used the function “glht” from the “multcomp” package (Hothorn et al., 2008) for Tukey post hoc tests of pair-wise differences in parasite numbers between colony types.

To analyse differences in parasite community composition, we used distance-based redundancy analyses with “management” or “colony type” as the constraining factor (dbRDA, function “db.rda” from the “vegan” package, (Oksanen et al., 2022)). Redundancy analysis summarizes multi-factor variation so that dissimilarities between samples can be graphically displayed in a two-dimensional coordinate system whose axes best separate the data based on predefined factors of interest (“constraints”). We chose to analyse parasite community compositions based on Jaccard distances (and thus based on presence/absence of parasite taxa) as opposed to Euclidean distance or Bray-Curtis dissimilarities (based on colony-level parasite abundance), because abundance data contained many zeroes and were non-normally distributed, and because it is hard to compare target molecule abundance variation of several orders of magnitude between parasites as different as arthropods and viruses. To partial out spatial structure, we conditioned our dbRDAs on principal coordinates of neighbour matrices (function “pcnm” from the “vegan” package; (Borcard & Legendre, 2002; Oksanen et al., 2022)). The model formulations were “db.rda(*Jaccard distance matrix of parasite communities* ~ X + Condition(scores(pcnm(*distance matrix of sample locations*)))”, with “X” denoting either “management” or “colony type”. This revealed that spatial location explained 17.7 % of the variation in parasite communities. Interestingly, a test with “study region” (four levels) as a constraining factor (formulation: “db.rda(*Jaccard distance matrix of parasite communities* ~ region)”) showed that 14.9 % of the variation could be attributed to differences between the four regions (see Fig. S1). This means that the spatial structure in parasite communities was mostly caused by large-scale (between regions) rather than fine-scale (between sites within one region) spatial variation. To infer the statistical significance of the constraining factors we used permutation tests with 99999 permutations (“anova.cca” function).

We did not perform separate statistical tests for each microparasite since potential interactions between taxa lead to non-independent data, and since the high number of individual tests would introduce statistical problems related to multiple testing. However, we graphically compared for each tested microparasite the prevalence and the associated 95% confidence intervals (Blaker, 2000; Stevenson et al., 2022) as well as the mean colony-level loads.

## Results

The number of microparasite taxa detected per colony ranged between 1 and 9, with an overall average of 5.8 taxa. Parasite counts differed significantly between regions (LRT: D.f.=6, χ^2^= 26.819, P=0.00016, Table S2); they were lowest in Coburg and Lichtenfels (mean: 4.9), intermediate on the Swabian Alb (mean: 5.7), and highest in Weilheim-Schongau (mean: 6.3) and Munich (mean: 6.2). On top of regional differences, management significantly affected the number of parasites per colony (LRT: D.f.=1, χ^2^= 14.677, P=0.00013, Table S3). Feral colonies had, on average, one parasite less (median: 5, mean: 5.4, range 1–8) than managed colonies (median: 6, mean: 6.2, range 4–9) (Fig. 2a). The difference between feral and managed colonies was consistent across study regions (no significant interaction between management and region; LRT: D.f.=3, χ^2^= 0.947, P=0.814, Table S4).

**Figure 2.**
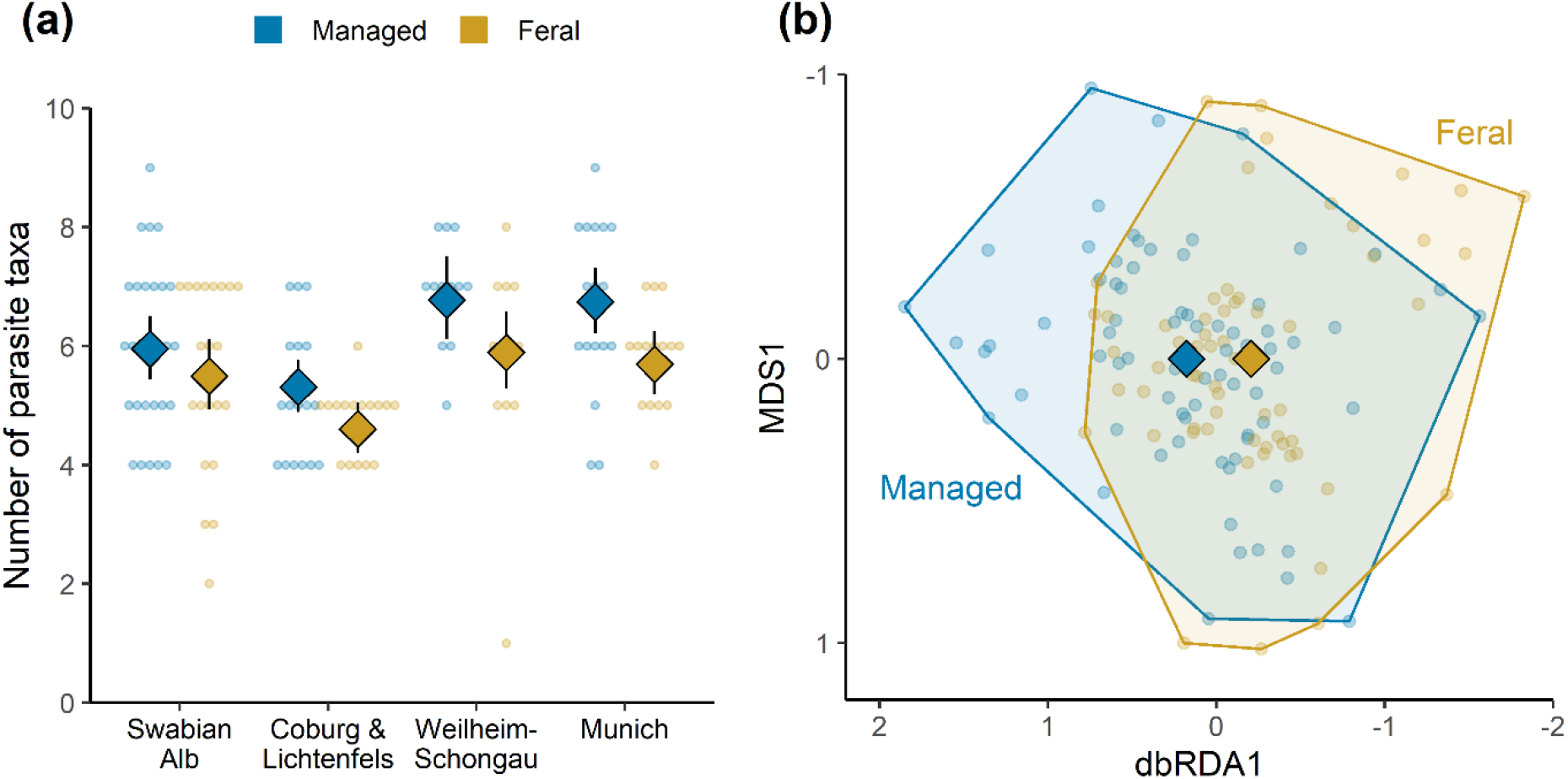
Comparison of parasite burden between managed (N=74) and feral (N=64) honeybee colonies. **(a)** Number of microparasite taxa detected among the 18 taxa assayed for each of the four study regions. Dots are raw data; diamond symbols and vertical lines give model-estimated means and 95% confidence intervals. See Table S5 for an overview of model predictions. **(b)** Representation of dissimilarities in microparasite communities as created by a distance-based redundancy analysis with management as the constraining factor (spatial structure partialled out). Managed and feral colonies are separated along the dbRDA1-axis (1.1% of total variation). The first unconstrained axis (MDS1) explains 16.7% of the variation. Dots represent locations of individual colonies and diamonds are mean locations.

The distance-based redundancy analysis revealed a marginal difference in the composition of parasite taxa between managed and feral colonies (permutational anova: P=0.074), with management explaining 1.1% of the variation in microparasite communities (Fig. 2b). A direct inspection of each of the 18 microparasites showed that prevalence was either similar in managed and feral colonies or lower in feral colonies (Fig. 3). A clear reduction in prevalence of more than 10 percentage points was found in four microparasite taxa: *Crithidia/Lotmaria* (Managed: 90.5%, Feral: 78.1%), Chronic bee paralysis virus (Managed: 14.9%, Feral: 1.6%), Deformed wing virus A (Managed: 18.9%, Feral: 3.1%), and Deformed wing virus B (Managed: 47.3%, Feral: 35.9%). Comparing mean colony-level parasite loads yielded a similar result, with abundances tending to be lower in feral colonies (Fig. 4a). Considering only colonies which were tested positive showed that parasite loads of infected feral and managed colonies were generally similar (Fig. 4b).

**Figure 3.**
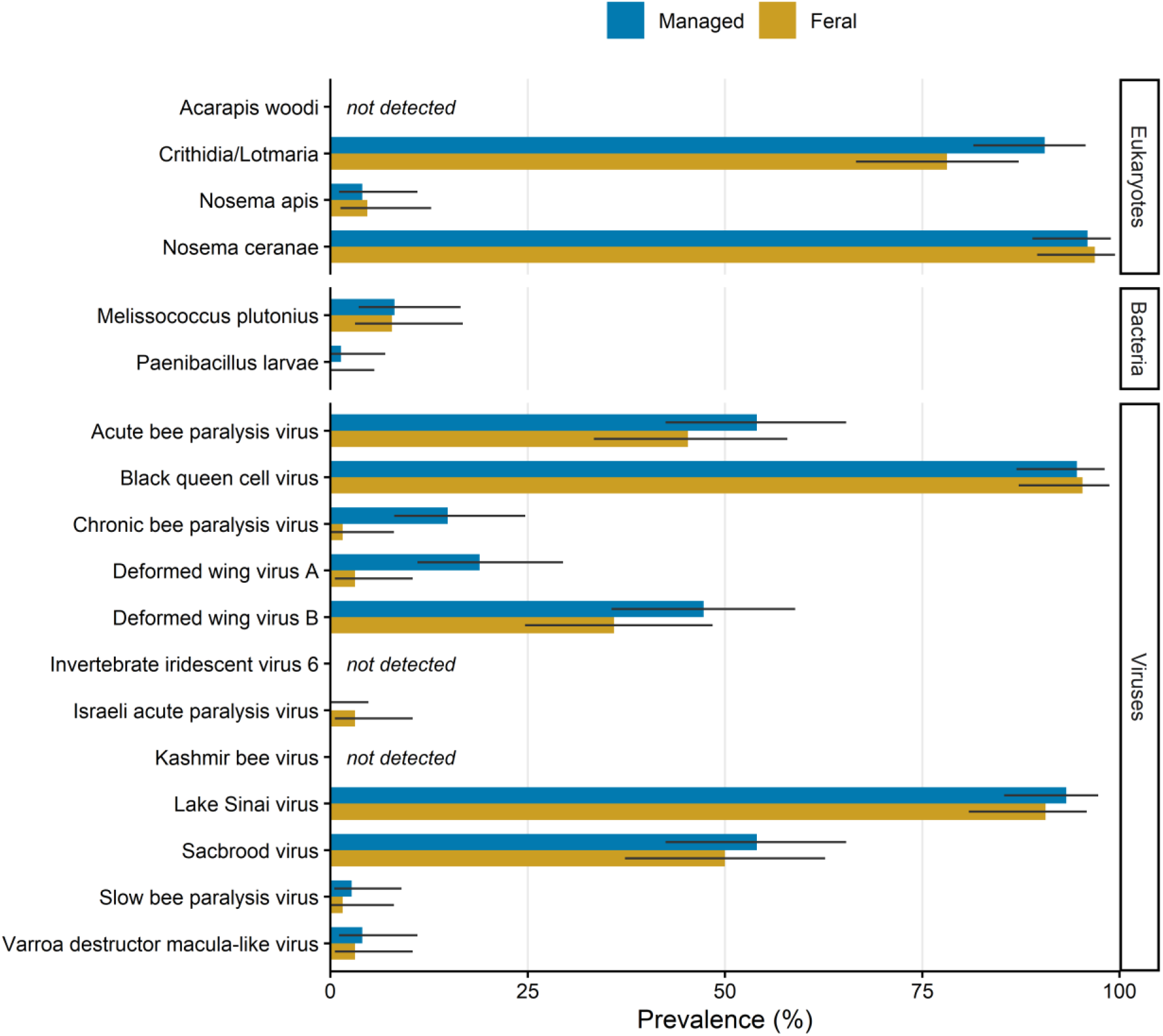
Prevalence (percentage of colonies tested positive; bars) and 95% confidence intervals (black lines) for 18 microparasite taxa among managed (N=74) and feral (N=64) honeybee colonies. See Table S6 for an overview of prevalence values and Tables S7–S10 for overviews of parasite prevalence divided by study region.

**Figure 4.**
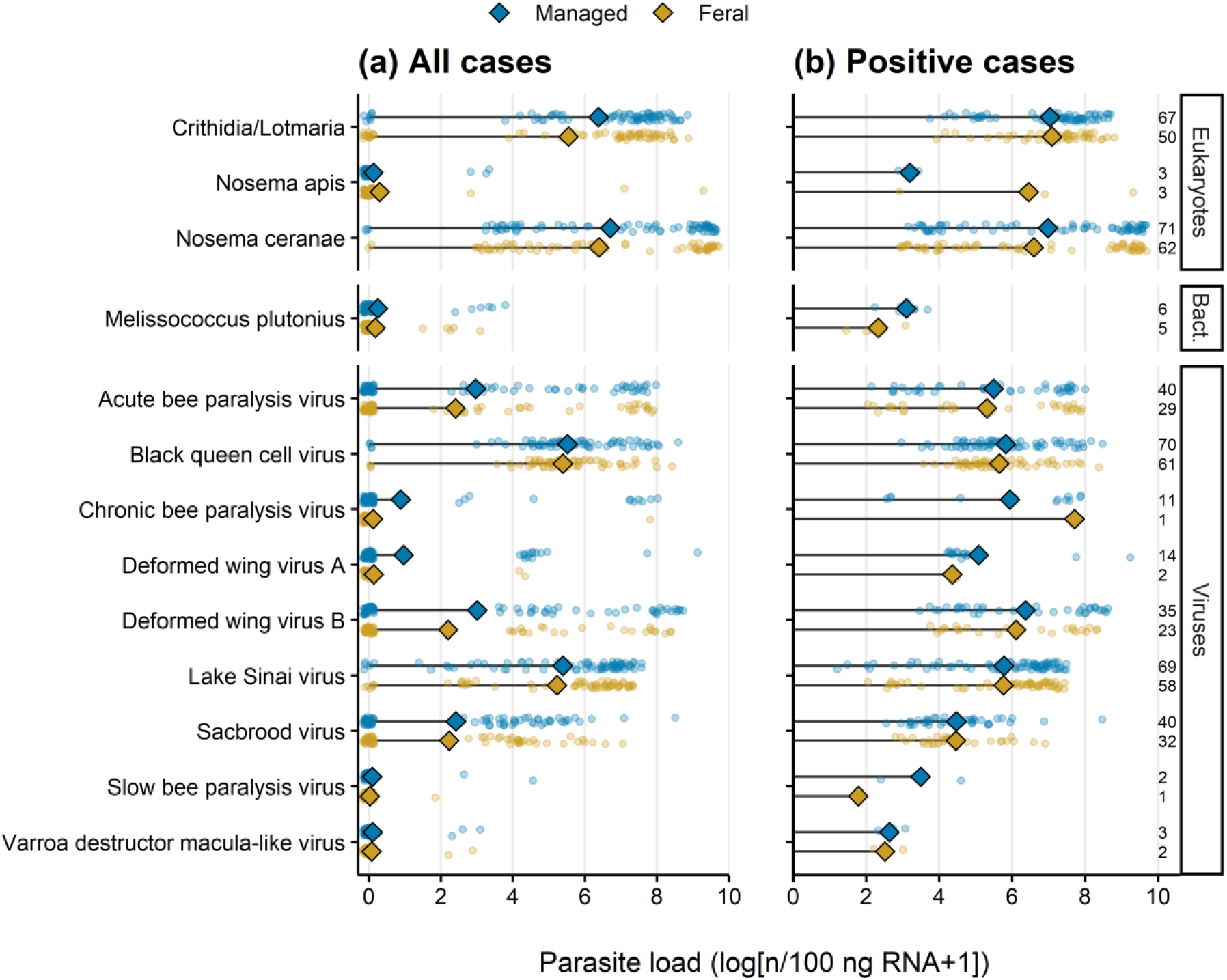
Parasite loads of managed and feral honeybee colonies. Only microparasite taxa detected at least once in each group are shown. **(a)** Parasite loads of individual colonies (dots; random jitter is added to reduce overplotting) and mean parasite loads (diamonds) based on all colonies tested (managed: N=74, feral: N=64). **(b)** Parasite loads based on positive cases only. Dots are parasite loads of individual colonies, diamonds are means, and numbers denote sample sizes. See Table S6 for an overview of parasite load values and Tables S7–S10 for overviews divided by study region.

An analysis of the number of parasites in relation to five colony types gave a more nuanced picture of differences in parasite burden (LRT, factor “colony type” after “region”: D.f.: 4, χ^2^=23.23, P=0.0001, Table S11). Parasite counts were lowest in newly founded feral colonies (mean: 5.3, range: 1–7), intermediate in overwintered feral colonies (mean: 5.7, range: 4–8) and nucleus colonies (mean: 5.7, range: 4–8), and highest in hived swarms (mean: 6.6, range: 4–8) and overwintered managed colonies (mean: 6.3, range: 4–9) (Fig. 5a). These differences were largely consistent across study regions (no significant interaction between colony type and region; LRT: D.f.: 12, χ^2^=9.546, P=0.656, Table S12; see Figure S2 and Table S13 for parasite counts divided by colony type and study region). A pairwise comparison revealed that the difference between feral founders and overwintered managed colonies (P<0.001) and between feral founders and hived swarms (P=0.002) were statistically significant. There were also differences in the colony-level composition of parasite taxa between the five colony types (colony type explained 3.3% of parasite community variation according to dbRDA; permutational anova: P=0.099; Fig. 5b). The differences in parasite communities, albeit marginal, resembled the differences in parasite counts: along the first dbRDA-axis, parasite assemblages of feral founders were separated from those of hived swarms and overwintered managed colonies, while the parasite communities of overwintered feral colonies took an intermediate position, and the parasite communities of managed nucleus colonies were closer to those of feral colonies than to those of other managed colonies (Fig. 5b). Accordingly, mean colony-level loads of individual parasite taxa tended to be lower in feral founders compared to overwintered feral colonies, and among the three types of managed colonies, mean parasite loads were most often lowest in nucleus colonies (Fig. 5c).

**Figure 5.**
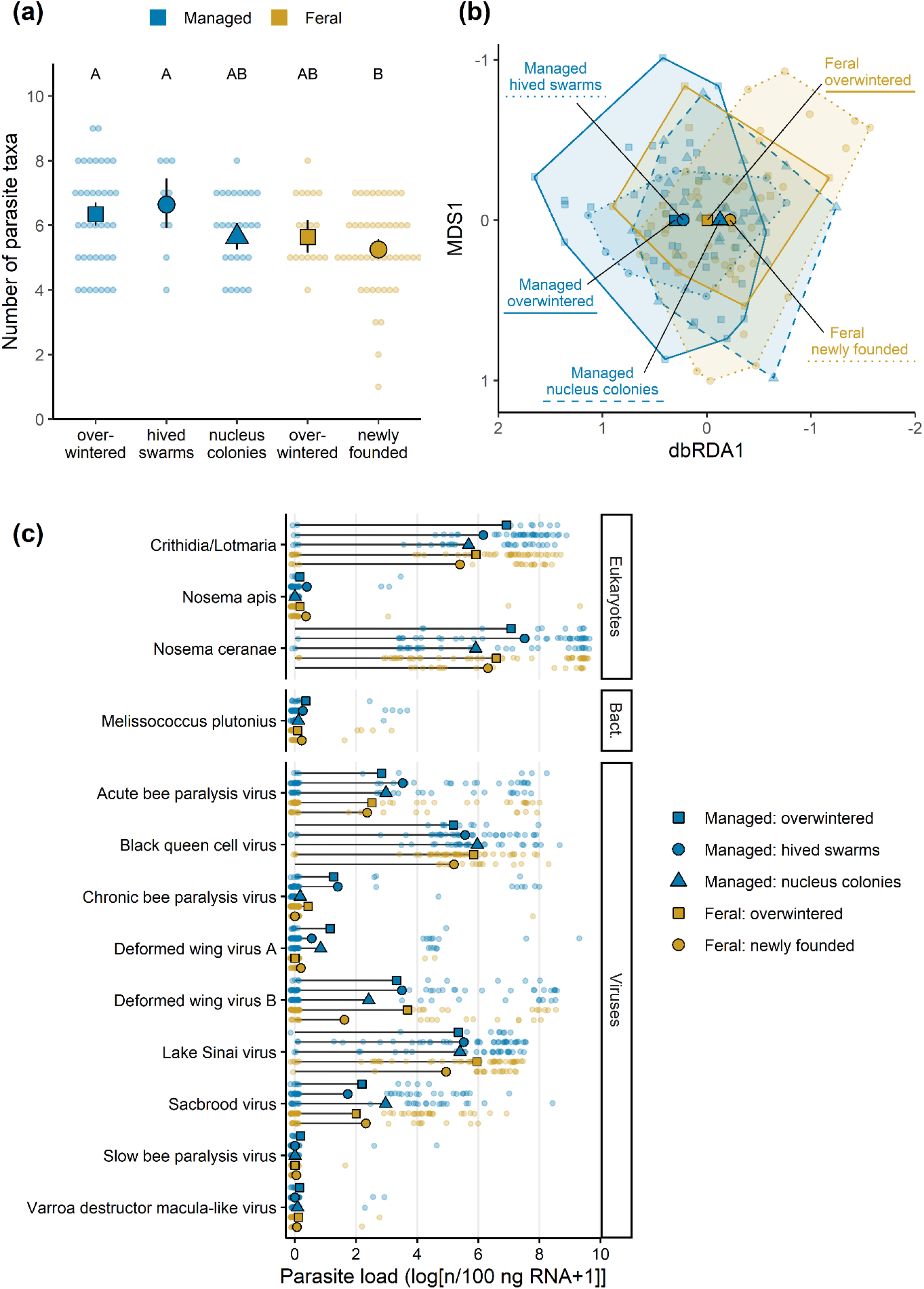
Comparison of parasite burden between different types of managed and feral honeybee colonies **(a)** Number of microparasite taxa detected among the 18 taxa assayed. Dots are raw data; large symbols and vertical lines give model-estimated means and 95% confidence intervals. Pairs that do not share a letter differ significantly (P<0.05). **(b)** Relative differences in microparasite community composition between the five colony types as revealed by a distance-based redundancy analysis with colony type as the constraining factor (spatial structure partialled out). The primary dbRDA-axis separating the colony types (1.9% of total variation) is shown along the first unconstrained axis (MDS1, 16.4% of the variation). Dots represent locations of individual colonies and diamonds are mean locations. **(c)** Parasite loads in relation to colony type. Dots are parasite loads of individual colonies and diamonds are means. Only microparasite taxa detected at least once among managed colonies and among feral colonies are shown (same selection as in Figure 4). See Table S14 for an overview of parasite prevalence and parasite loads by colony type.

## Discussion

We compared the parasite burden of honeybee colonies in managed and feral conditions to evaluate whether colonies that escaped from apiaries pose a risk to managed and wild bee health by acting as reservoirs of disease-causing agents. The number of microparasites detected per colony and the prevalence of four important taxa were clearly lower in feral compared to managed colonies. This was explained by differences in population demography, with most feral colonies being recent founders with very low parasite loads, and by environmental differences between managed and feral colonies. We conclude that feral honeybee colonies are unlikely to contribute disproportionately to the spread of bee parasites.

We considered parasite burden based on the prevalence of 18 microparasites determined using qPCR. This means that we did not cover all known bee parasites. We also cannot exclude the possibility of some false non-detections of targeted RNA viruses due to their high evolutionary rates. However, our conclusions about relative differences in parasite burden between groups are robust because all colonies were subjected to the same qPCR-assays and because groups were evenly distributed across study regions.

Comparing colonies from four regions in southern Germany revealed strong geographic differences in parasite numbers and parasite community composition. These differences might be explained by a combination of factors including managed colony density, land use, climate, and colonization histories of individual parasites. The management implication is that moving managed hives, even over moderate distances (the distance between our study regions Weilheim-Schongau and Munich is approx. 50 km), bears the risk of introducing bee parasites that would otherwise not be present locally. Considering the potential negative effects on local wild bee communities and on honeybees managed by non-migratory beekeepers (Martínez-López et al., 2022), apicultural disease management needs to pay more attention to regional differences in parasite communities.

Although region of sampling had a stronger effect on parasite communities than management, we believe that the differences found between feral and managed colonies are ecologically relevant. With about six parasite taxa detected per colony on the overall average, the reduced count of one parasite per colony from managed to feral colonies is noteworthy, especially since this pattern was consistent across study regions. Higher numbers of parasites have been linked to higher colony mortality (vanEngelsdorp et al., 2009), although this relationship might be non-linear (Ravoet et al., 2013). Analysing the associations between 18 parasite taxa and colony mortality, Ravoet et al. (2013) found that winter mortality steeply increased with the number of parasite species rising from 3 to 5, but additional parasites had no further effect. In our study, the percentage of colonies with more than 5 detected parasite taxa was only 47 % in the group of feral colonies but 65 % in the group of managed colonies, hence the difference was likely within a relevant range.

Importantly, it was not entirely random which parasites were less prevalent in feral colonies: Trypanosomatidae, chronic bee paralysis virus and deformed wing virus strains A and B were less frequently detected in feral colonies. These four parasites are all important since they can induce host mortality. Furthermore, they have been detected in other bee species, so they are also relevant in the context of honeybee-wild bee interactions (Strobl et al., 2019; Tehel et al., 2016). The two trypanosomatid species *Chrithidia mellificae* and *Lotmaria passim* (among which we did not distinguish with our PCR assay) are unicellular parasites that colonize the bees’ hindgut (Schwarz et al., 2015). They have been experimentally shown to reduce the lifespan of individual workers (Strobl et al., 2019) and their presence is associated with colony-level winter mortality (Ravoet et al., 2013). Chronic bee paralysis virus (CBPV) causes the bee paralysis disease in adult workers, which involves symptoms like hair loss, undirected trembling walks, and loss of flight ability. The virus can spread in a colony via worker contact and eventually lead to its collapse (Ribière et al., 2010). Interestingly, CBPV seems to be an emerging threat, as indicated by a rapid rise of cases among Britain’s apiaries during the last decade (Budge et al., 2020). Given that in this study, its prevalence was 14.9% in managed colonies but only 1.6% in feral colonies, the latter are unlikely to represent an important dispersal route for the virus.

The reduction in the prevalence of deformed wing viruses (DWV) A and B in feral compared to managed colonies is, ecologically, the most important detected difference since DWV is regarded as one of the main causes of winter colony losses (Dainat et al., 2012a; de Miranda & Genersch, 2010; Genersch et al., 2010). When transmitted during development, DWV directly kill their host at the pupal stage or cause body deformations of the emerging bee, leading to premature death (de Miranda & Genersch, 2010). Even infected bees that do not show overt symptoms have their life expectancy reduced by DWV, which can result in mass losses of bees on the colony level and its collapse during winter (Dainat et al., 2012a). Importantly, the abundance of DWV in honeybee colonies and the severity of the resulting symptoms are tightly linked to the co-occurrence with *V. destructor* (de Miranda & Genersch, 2010; Paxton et al., 2022). While DWV were unproblematic before the invasion of *V. destructor*, the mite represented a new vector not only aiding the spread but also fuelling the evolution of the viruses, as the emergence and ongoing replacement of DWV-A by the presumably more virulent DWV-B indicates (McMahon et al., 2016; Paxton et al., 2022). In turn, the mite’s reproduction is enhanced by the presence of DWV, making the mite-virus pair a deadly symbiosis (Di et al., 2016). Unfortunately, it was not possible in our study to investigate the levels of *V. destructor* infestation, because the numbers of mites on foragers are generally too low to make a reasonable analysis based on bees captured at the colony entrance. It would have required sampling pupae or young bees from the brood nest or assessing the rate at which dead mites naturally fall to the bottom of the nest cavity (Dietemann et al., 2013), but this was not possible in the case of the feral colonies. However, since DWV and *Varroa* correlate (Dainat et al., 2012b; Norton et al., 2021), the lower prevalence of DWV in feral colonies suggests that mite infestation levels were also reduced.

Our findings are in seeming conflict with a previous study from England which concluded that parasite pressure on feral colonies is relatively high. Thompson et al. (2014) tested for 10 parasites and found that these were equally prevalent among managed and feral colonies, but the colony-level parasite abundance of one parasite, deformed wing virus, was significantly higher in feral colonies (no difference was made between DWV-A and DWV-B). However, the sample of feral colonies analysed in their study only included colonies aged at least one year. We also found that the parasite prevalence in feral colonies aged at least one year (overwintered feral colonies) did not differ significantly from managed colonies and that they had relatively high loads of DWV-B (although not of DWV-A; Fig. 5c). Therefore, had we only considered old feral colonies, we might have come to similar conclusions. However, we would have then seriously overestimated the parasite burden in the feral population as a whole because only about 10% of the feral colonies present in summer are older than one year (Kohl et al., 2022), and especially the young, newly founded feral colonies had their parasite burden reduced compared to managed colonies. These considerations demonstrate that it is important to know the population demography of the feral honeybees under consideration when asking questions about their ecological impact, e.g., their contribution to the spread of bee diseases.

The feral honeybees investigated in this study are known to be recent descendants from colonies managed in apiaries (Kohl et al., 2022), and therefore, their reduced parasite burden needs to be explained by ecological/environmental, rather than genetic, differences from managed honeybees. By considering five colony types representing differences in colony age and management history (Table 1), we gained some insights into why parasite burden is reduced in feral colonies. We found that newly founded feral colonies had a lower parasite burden than both overwintered feral colonies and overwintered managed colonies (from which many of the young feral colonies directly descend). This supports the hypothesis that the natural process of reproductive swarming – the abandonment of the old nest, the pause of brood production, and the construction of fresh comb at a new dwelling place – leads to a temporal release from parasite pressure (DeGrandi-Hoffman et al., 2017; Loftus et al., 2016; Royce et al., 1991). The high proportion of young, swarm-founded colonies in the feral population is one of the reasons for the overall significant difference in parasite burden between feral and managed colonies. In the managed population, the proportion of young colonies is much lower (about 50 %), and most young colonies are so-called nucleus colonies, created by transferring several combs with brood from the mother colony into a new hive – a completely different founding mechanism. Since nucleus colonies directly inherit the parasites residing in the old combs and the brood, their parasite communities should resemble those of established (old) colonies. Indeed, parasite counts did not differ significantly between nucleus colonies and overwintered managed colonies, nor between nucleus colonies and overwintered feral colonies, albeit overwintered managed colonies had the highest numbers and a different community composition of parasites. The latter might be explained by the fact that beekeepers usually manage established colonies in such a way that brood is continuously produced, while managed nucleus colonies and overwintered feral colonies typically experience a brood pause in spring and thus a temporal reduction of the breeding ground for parasites (Table 1) (Loftus et al., 2016). Importantly, young managed colonies founded by swarms (“hived swarms”) were infested by a significantly higher number of microparasite taxa than young feral colonies. This contrast needs to be regarded with care since our sample size of hived swarms was low (N=9), but it suggests that not only differences in population age structure between managed and feral colonies, but also in the environment, contribute to the difference in parasite burden.

**Table 1:**
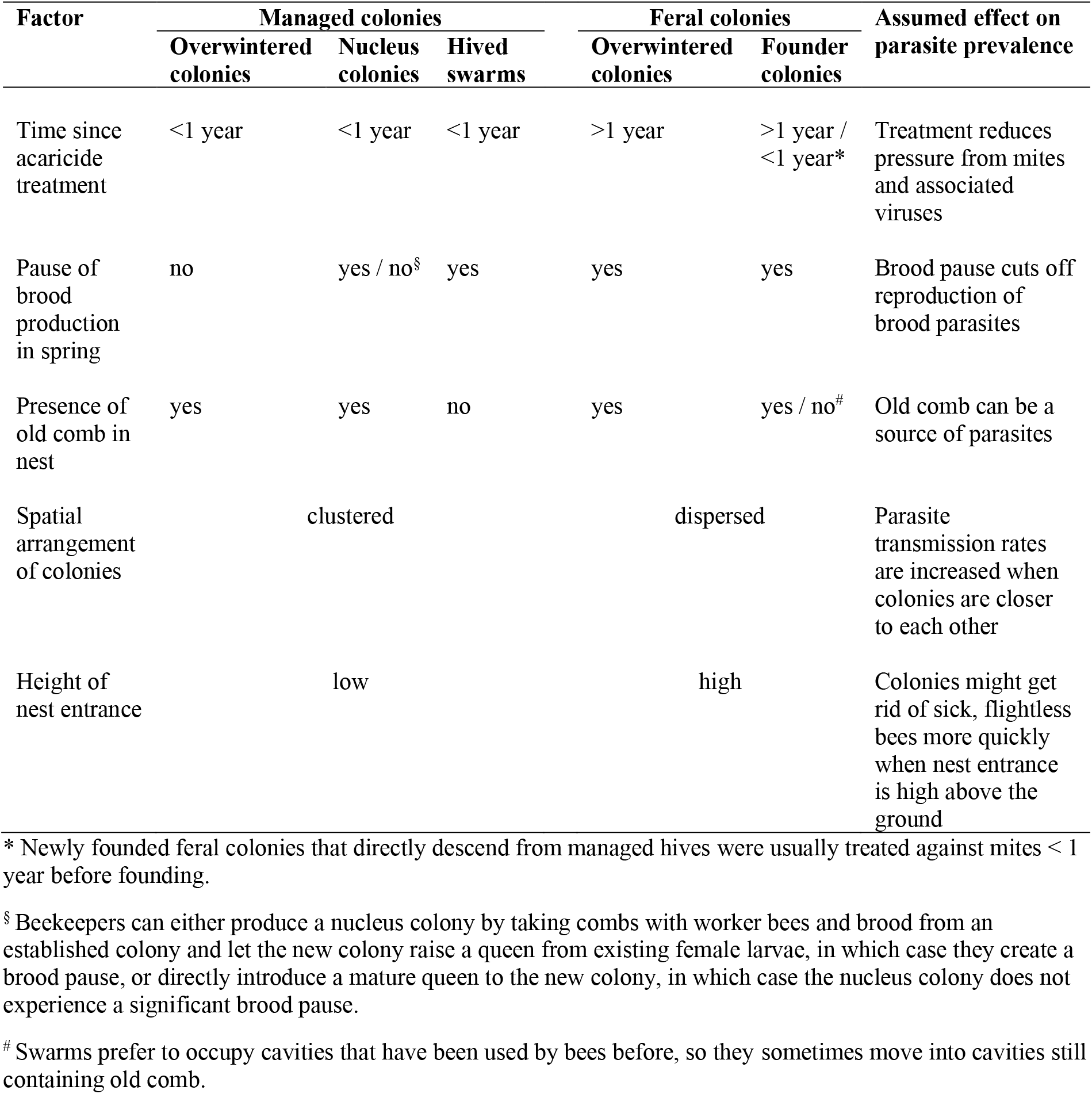
Nonexclusive list of factors that might affect parasite burden, and their typical manifestation in different types of managed and feral honeybee colonies.

The most obvious environmental difference that has been demonstrated to affect parasite pressure is that managed hives are typically clustered in apiaries, while feral colonies are spatially dispersed (Lindström et al., 2008; Nolan & Delaplane, 2017; Seeley & Smith, 2015). The crowding of colonies promotes random drifting and robbing behaviour, and both drifters and robbers can transfer parasites between hives (Lindström et al., 2008; Peck & Seeley, 2019; Seeley & Smith, 2015). Another highly consistent difference is that beekeepers conventionally keep their hives at ground level, while feral colonies typically nest in cavities several meters above the ground. The bees’ preference for aerial cavities is thought to serve as a protection against predators, but a positive side effect might be that colonies dispose of sick bees more easily. For example, both CBPV and DWV cause the loss of flight ability, and flightless bees that fall out of the nest might struggle to find their way back when the entrance is high above the ground. There are more differences which might directly or indirectly affect the colonies’ likelihood of becoming infected by parasites and of developing disease symptoms, e.g., the size of the nest entrance, the presence of landing boards to aid bees entering the hive, the insulation of the hive, the microclimate at the nest site, the frequency of disturbance, or the type of food consumed. Given the role parasites play in limiting the survival of managed colonies in apiculture, it seems worth experimentally investigating these factors in more detail. For example, an interesting new question raised by our study is whether it is possible to restore the natural parasite-reducing effect of swarm-founding under apicultural management.

Feral honeybee colonies carrying many parasites and/or high parasite abundances would be a nuisance to apicultural disease management and would pose a risk to the health of *non-Apis* wild bees. However, we found that feral colonies have on average fewer parasites and lower parasite loads than managed colonies. This is partly explained by the effect of natural swarm reproduction and dispersal – a mechanism of parasite reduction that might also be effective in other swarm-founding social insects (McGlynn, 2012). Given that feral colonies have a relatively low parasite burden and that they make up a small fraction of the overall honeybee population (in Germany feral colonies make up about 5% of the whole honeybee population in summer), we conclude that it is unlikely that they significantly contribute to the spread of bee parasites. On the contrary, new disease agents are probably primarily propagated by managed colonies, as indicated by the higher prevalence of two emerging viruses, CBPV and DWV-B, in the sample of managed hives. The management implication of this work is that the prevention of epidemics is no suitable argument for the often-practised removal or destruction of feral honeybee nests (Taylor et al., 2007). In fact, our data suggest that there is no conflict between the promotion of wild-living honeybee populations and the management of bee diseases in apiculture. What remains unclear is how the various environmental differences between wild nests and hives at apiaries contribute to the reduced parasite burden in feral colonies. Some known natural parasite-reducing factors, e.g., the spatial separation of colonies and the periodic interruption of brood production, can readily be adopted by beekeepers to increase the health of managed honeybees and to reduce the risk of disease spread by apiculture (Büchler et al., 2020; Dynes et al., 2019; Loftus et al., 2016).

## Supporting information

Supplementary information for Reduced parasite burden in feral honeybee colonies

## Authors’ contributions

P.L.K.: conceptualization, formal analysis, funding acquisition, investigation, methodology, visualization, writing–original draft, writing–review & editing; P.D..: formal analysis, investigation, methodology, validation, writing–review & editing; B.R.: conceptualization, investigation, methodology, writing–review & editing; S.R.: conceptualization, investigation, methodology, writing–review & editing; F.R.: conceptualization, investigation, methodology, writing–review & editing; I.S.D.: conceptualization, supervision, writing–review & editing; M.H.: resources, supervision, writing–review & editing.

## Acknowledgements

We thank Luis Sikora, Norbert Wimmer and Kurt Zeimentz for sharing locations of woodpecker cavities which serve as nest sites for feral honeybees in German forests. We also thank all beekeepers who granted us permission to sample bees from their hives. This study was supported German Federal Environmental Foundation (DBU).

