## Supplementary information for Reduced parasite burden in feral honeybee colonies for "Reduced parasite burden in feral honeybee colonies"

by

Patrick L. Kohl*, Paul D'Alvise, Benjamin Rutschmann, Sebastian Roth, Felix Remter,

Ingolf Steffan-Dewenter & Martin Hasselmann

### RNA extraction and qPCR analyses

We obtained colony-level total RNA by extracting it from multi-bee homogenates using a TRIzol protocol. For each colony, 20 workers were placed in a 15 mL reaction tube with one 0.25-inch ceramic bead (MP Biomedicals), five 2.8-mm Precellys® steal beads, 0.7 g 0.1-mm glass/zirconia beads (BioSpec) and 4 mL TRIzol (Invitrogen). The mixtures were homogenized with a FastPrep24 (MP Biomedicals) running two times at 6 m/s speed for 60 s (in between the runs, tubes were inverted and vigorously shaken by hand to remove bees stuck to the bottom of the tubes). The homogenates were incubated for five minutes at room temperature (RT), mixed with 800 µL chloroform, vigorously shaken for 15 s, and incubated for another five minutes at RT. After 15 min of centrifugation at 12,000 g and 4°C, 200 µL of the aqueous phases were transferred to 1.5-mL reaction tubes and mixed with 250 µL isopropanol by repeated inverting. After another incubation for 10 min at RT, the precipitated RNA was separated by centrifugation (12,000 g, 4°C). The resulting supernatants were removed, and the RNA pellets were washed with 75% ethanol, dried for 5 min at RT, and redissolved in 50 µL nuclease-free water. RNA concentrations (determined using a NanoDrop spectrophotometer, Themo Fisher) ranged between 2215 and 6117 ng/µL (mean: 3519 ng/µL). They were used as references for calculating colony-level parasite loads. Production of cDNA, pre-amplification of target sequences, qPCR on a Biomark HD system, and calculation of number of target molecules from Cq-values were performed as described in D'Alvise et al. (2019).

**Table S1**: Overview of the primers used in this study.

| PCR target | Primers 5'–3' | Reference |
|---|---|---|
| *Acarapis woodi* | F: GGAATATGATCTGGTTTAGTTGGTC<br>R: GAATCAATTTCCAAACCCACCAATC | Cepero et al., 2015 |
| *Crithidia/Lotmaria* | F: CCGCTTTTGGTCGGTGGAGTGAT<br>R: GCAGGGACGTAATCGGCACAGTTT | D'Alvise et al., 2019 |
| *Nosema apis* | F: CAGTTATGGGAAGTAACATAGTTG<br>R: CGATTTGCCCTCCAATTAATCTG | D'Alvise et al., 2019 |
| *Nosema ceranae* | F: TGAGGCAGTTATGGGAAGTAATATTATATTG<br>R: ACTTGATTTGCCCTCCAATTAATCAC | D'Alvise et al., 2019 |
| **Bacteria** | | |
| *Melissococcus plutonius* | F: TGTTGTTAGAGAAGAATAGGGGAA<br>R: CGTGGCTTTCTGGTTAGA | Budge et al., 2010 |
| *Paenibacillus larvae* | F: CGGGAGACGCCAGGTTAG<br>R: TTCTTCCTTGGCAACAGAGC | Martínez et al., 2010 |
| **Viruses** | | |
| Acute bee paralysis virus | F: TCATACCTGCCGATCAAG<br>R: CTGAATAATACTGTGCGTATC | Locke et al., 2012 |
| Black queen cell virus | F: AGTGGCGGAGATGTATGC<br>R: GGAGGTGAAGTGGCTATATC | Locke et al., 2012 |
| Chronic bee paralysis virus | F: CAACCTGCCTCAACACAG<br>R: AATCTGGCAAGGTTGACTGG | Locke et al., 2012 |
| Deformed wing virus A | F: TTCATTAAAGCCACCTGGAACATC<br>R: TTTCCTCATTAACTGTGTCGTTGA | Locke et al., 2012 |
| Deformed wing virus B | F: GCCCTGTTCAAGAACATG<br>R: CTTTTCTAATTCAACTTCACC | Locke et al., 2012 |
| Invertebrate iridescent virus 6 | F: TGGTTYACCCAAGTACCKGTTAG<br>R: ATGCKGACCATTCGCTTC | Papp et al., 2014 |
| Israeli acute paralysis virus | F: CCATGCCTGGCGATTCAC<br>R: CTGAATAATACTGTGCGTATC | Locke et al., 2012 |
| Kashmir bee virus | F: CCATACCTGCTGATAACC<br>R: CTGAATAATACTGTGCGTATC | Locke et al., 2012 |
| Lake Sinai virus | F: TCATCCCAAGAGAACCAC<br>R: GCATGGAAGAGAGTAGGTA | D'Alvise et al., 2019 |
| Sacbrood virus | F: TTGGAACTACGCATTCTCTG<br>R: GCTCTAACCTCGCATCAAC | Locke et al., 2012 |
| Slow bee paralysis virus | F: GCGCTTTAGTTCAATTGCC<br>R: ATTATAGGACGTGAAAATATAC | Locke et al., 2012 |
| Varroa destructor macula-like virus | F: ATCCCTTTTCAGTTCGCT<br>R: AGAAGAGACTTCAAGGAC | Locke et al., 2012 |
| **Control genes** | | |
| Actin | F: TGCCAACACTGTCCTTTCTG<br>R: AGAATTGACCCACCAATCCA | Lourenço et al., 2008 |
| Elongation factor 1 | F: GGAGATGCTGCCATCGTTAT<br>R: CAGCAGCGTCCTTGAAAGTT | Lourenço et al., 2008 |
| Ribosomal protein S5 | F: AATTATTTGGTCGCTGGAATTG<br>R: TAACGTCCAGCAGAATGTGGTA | Evans, 2006 |

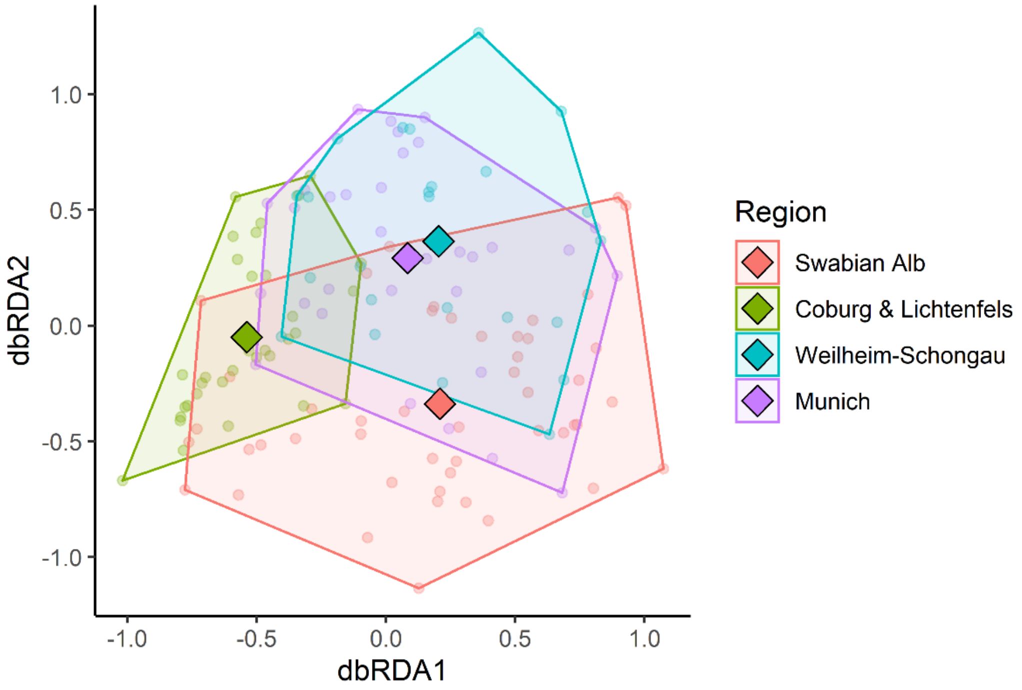


**Figure S1**: Dissimilarity of microparasite communities in honeybee colonies (N=138) between the four study regions as revealed by a redundancy analysis with “region” as a constraining factor. Region of sampling explains 14.9% of variation in parasite community composition based on presence/absence of taxa (Jaccard distance). The first two constrained axes explain 6.5 % (dbRDA1) and 5.2% (dbRDA2) of the variation. Dots are colony locations and diamonds are mean locations.

**Table S2**: Result of a likelihood ratio test (function "anova()" in R) comparing two nested models of the number of detected parasites. The factor "region" significantly improves model fit compared to a null model.

| Model | Df | AIC | BIC | logLik | deviance | Chisq | Δ Df | Pr(>Chisq) |
|---|---|---|---|---|---|---|---|---|
| glmmTMB(Number of parasites **~ 1**, family = genpois()) | 2 | 487.199 | 493.054 | -241.600 | 483.199 | | | |
| glmmTMB(Number of parasites **~ region**, dispformula = ~region, family = genpois()) | 8 | 472.380 | 495.798 | -228.190 | 456.380 | 26.819 | 6 | 0.00016 |

**Table S3**: Result of a likelihood ratio test (function "anova()" in R) comparing two nested models explaining the number of detected parasites. Adding the factor "management" (feral versus managed) to the factor "region" significantly improves model fit.

| Model | Df | AIC | BIC | logLik | deviance | Chisq | Δ Df | Pr(>Chisq) |
|---|---|---|---|---|---|---|---|---|
| glmmTMB(Number of parasites **~ region,** dispformula = ~region, family = genpois()) | 8 | 472.380 | 495.798 | -228.190 | 456.380 | | | |
| glmmTMB(Number of parasites ~ **management + region**, dispformula = ~region, family = genpois()) | 9 | 459.703 | 486.048 | -220.852 | 441.703 | 14.677 | 1 | 0.00013 |

**Table S4**: Result of a likelihood ratio test (function "anova()" in R) comparing two nested models of the number of detected parasites. The interaction between management and region does not significantly improve model fit.

| Model | Df | AIC | BIC | logLik | deviance | Chisq | Δ Df | Pr(>Chisq) |
|---|---|---|---|---|---|---|---|---|
| glmmTMB(Number of parasites ~ **management + region**, dispformula = ~region, family = genpois()) | 9 | 459.703 | 486.048 | -220.852 | 441.703 | | | |
| glmmTMB(Number of parasites ~ **management + region + management:region**, dispformula= ~region, family = genpois()) | 12 | 464.756 | 499.883 | -220.378 | 440.756 | 0.947 | 3 | 0.814 |

**Table S5:** Mean numbers of detected parasites per colony and the respective 95% confidence limits (CI) estimated by a generalized linear model (model formula: “glmmTMB(Number of parasites ~ management + region + management : region, dispformula= ~ region, family = genpois()”) for all cases (overall), for managed cases and for feral cases, and divided by study region.

| **Region** | **Management** | **Estimate** | **Lower CI** | **Upper CI** |
|---|---|---|---|---|
| *Overall* | *Overall* | 5.768 | 5.573 | 5.97 |
| | Managed | 6.164 | 5.895 | 6.446 |
| | Feral | 5.397 | 5.132 | 5.676 |
| Swabian Alb | *Overall* | 5.718 | 5.329 | 6.135 |
| | Managed | 5.951 | 5.444 | 6.506 |
| | Feral | 5.493 | 4.938 | 6.111 |
| Coburg & Lichtenfels | *Overall* | 4.946 | 4.642 | 5.271 |
| | Managed | 5.951 | 5.444 | 6.506 |
| | Feral | 4.605 | 4.204 | 5.045 |
| Weilheim-Schongau | *Overall* | 6.318 | 5.853 | 6.820 |
| | Managed | 6.775 | 6.113 | 7.508 |
| | Feral | 5.892 | 5.278 | 6.576 |
| Munich | *Overall* | 6.195 | 5.815 | 6.601 |
| | Managed | 6.741 | 6.214 | 7.313 |
| | Feral | 5.693 | 5.184 | 6.253 |

**Table S6:** Overview of prevalence and colony-level loads of 15 detected microparasites for A: all colonies tested, M: managed colonies only, and F: feral colonies only. Three other parasites assayed in this study (*Acarapis woodi*, Invertebrate iridescent virus 6, and Kashmir bee virus) were not detected in any colony.

| | | | | Prevalence (%) | | | $Log_{10}$ (n / 100 ng RNA+1) | | |
|---|---|---|---|---|---|---|---|---|---|
| **Parasite Taxon** | **Sample** | **N** | **N (positives)** | **Estimate** | **Lower CI** | **Upper CI** | **Mean** | **Mean (positives)** | **Max** |
| *Crithidia/ Lotmaria* | A | 138 | 117 | 84.8 | 77.7 | 90.1 | 5.99 | 7.06 | 8.84 |
| | M | 74 | 67 | 90.5 | 81.5 | 95.7 | 6.37 | 7.04 | 8.75 |
| | F | 64 | 50 | 78.1 | 66.6 | 87.2 | 5.54 | 7.09 | 8.84 |
| *Nosema apis* | A | 138 | 6 | 4.3 | 1.9 | 9.2 | 0.21 | 4.82 | 9.37 |
| | M | 74 | 3 | 4.1 | 1.1 | 11 | 0.13 | 3.19 | 3.49 |
| | F | 64 | 3 | 4.7 | 1.3 | 12.8 | 0.3 | 6.45 | 9.37 |
| *Nosema ceranae* | A | 138 | 133 | 96.4 | 91.9 | 98.6 | 6.55 | 6.80 | 9.59 |
| | M | 74 | 71 | 95.9 | 89 | 98.9 | 6.7 | 6.98 | 9.57 |
| | F | 64 | 62 | 96.9 | 89.6 | 99.4 | 6.39 | 6.593 | 9.59 |
| **Bacteria** | | | | | | | | | |
| *Melissococcus plutonius* | A | 138 | 11 | 8 | 4.2 | 13.6 | 0.22 | 2.75 | 3.71 |
| | M | 74 | 6 | 8.1 | 3.6 | 16.5 | 0.25 | 3.10 | 3.71 |
| | F | 64 | 5 | 7.8 | 3.1 | 16.8 | 0.18 | 2.32 | 3.22 |
| *Paenibacillus larvae* | A | 138 | 1 | 0.7 | 0 | 3.7 | 0.03 | 3.68 | 3.68 |
| | M | 74 | 1 | 1.4 | 0.1 | 7 | 0.05 | 3.68 | 3.68 |
| | F | 64 | 0 | 0 | 0 | 5.6 | 0 | NA | 0 |
| **Viruses** | | | | | | | | | |
| Acute bee paralysis virus | A | 138 | 69 | 50 | 41.6 | 58.4 | 2.71 | 5.42 | 8.12 |
| | M | 74 | 40 | 54.1 | 42.5 | 65.3 | 2.97 | 5.49 | 8.12 |
| | F | 64 | 29 | 45.3 | 33.4 | 57.9 | 2.41 | 5.31 | 7.90 |
| Black queen cell virus | A | 138 | 131 | 94.9 | 90.1 | 97.7 | 5.45 | 5.74 | 8.55 |
| | M | 74 | 70 | 94.6 | 86.9 | 98.1 | 5.51 | 5.83 | 8.55 |
| | F | 64 | 61 | 95.3 | 87.2 | 98.7 | 5.38 | 5.65 | 8.30 |
| Chronic bee paralysis virus | A | 138 | 12 | 8.7 | 4.8 | 14.6 | 0.53 | 6.09 | 7.93 |
| | M | 74 | 11 | 14.9 | 8.1 | 24.7 | 0.88 | 5.94 | 7.93 |
| | F | 64 | 1 | 1.6 | 0.1 | 8 | 0.12 | 7.71 | 7.71 |
| Deformed wing virus A | A | 138 | 16 | 11.6 | 7 | 17.9 | 0.58 | 4.99 | 9.28 |
| | M | 74 | 14 | 18.9 | 11 | 29.5 | 0.96 | 5.08 | 9.28 |
| | F | 64 | 2 | 3.1 | 0.6 | 10.4 | 0.14 | 4.36 | 4.45 |
| Deformed wing virus B | A | 138 | 58 | 42 | 34 | 50.6 | 2.63 | 6.27 | 8.63 |
| | M | 74 | 35 | 47.3 | 35.6 | 58.9 | 3.01 | 6.37 | 8.63 |
| | F | 64 | 23 | 35.9 | 24.6 | 48.4 | 2.2 | 6.11 | 8.41 |
| Israeli acute paralysis virus | A | 138 | 2 | 1.4 | 0.3 | 5.1 | 0.02 | 1.34 | 1.45 |
| | M | 74 | 0 | 0 | 0 | 4.8 | 0 | NA | 0 |
| | F | 64 | 2 | 3.1 | 0.6 | 10.4 | 0.04 | 1.34 | 1.45 |
| Lake Sinai virus | A | 138 | 127 | 92 | 86.4 | 95.8 | 5.31 | 5.77 | 7.58 |
| | M | 74 | 69 | 93.2 | 85.4 | 97.3 | 5.38 | 5.78 | 7.58 |
| | F | 64 | 58 | 90.6 | 80.9 | 95.8 | 5.22 | 5.77 | 7.41 |
| Sacbrood virus | A | 138 | 72 | 52.2 | 43.8 | 60.6 | 2.33 | 4.46 | 8.37 |
| | M | 74 | 40 | 54.1 | 42.5 | 65.3 | 2.42 | 4.47 | 8.37 |
| | F | 64 | 32 | 50 | 37.3 | 62.7 | 2.23 | 4.46 | 6.98 |
| Slow bee paralysis virus | A | 138 | 3 | 2.2 | 0.6 | 6.2 | 0.06 | 2.92 | 4.49 |
| | M | 74 | 2 | 2.7 | 0.5 | 9 | 0.09 | 3.49 | 4.49 |
| | F | 64 | 1 | 1.6 | 0.1 | 8 | 0.03 | 1.78 | 1.78 |
| Varroa destructor macula-like virus | A | 138 | 5 | 3.6 | 1.4 | 8.1 | 0.09 | 2.58 | 3.02 |
| | M | 74 | 3 | 4.1 | 1.1 | 11 | 0.11 | 2.63 | 3.02 |
| | F | 64 | 2 | 3.1 | 0.6 | 10.4 | 0.08 | 2.51 | 2.91 |

**Table S7:** Overview of prevalence and colony-level loads of 15 microparasites for study region 1 (Swabian Alb). Three other parasites assayed in this study (*Acarapis woodi*, Invertebrate iridescent virus 6, and Kashmir bee virus) were not detected any of the four study regions. A: all colonies, M: managed colonies, and F: feral colonies.

| | | | | **Prevalence (%)** | | | **$Log_{10}$ (n / 100 ng RNA+1)** | | |
|---|---|---|---|---|---|---|---|---|---|
| **Parasite Taxon** | **Sample** | **N** | **N (positives)** | **Estimate** | **Lower CI** | **Upper CI** | **Mean** | **Mean (positives)** | **Max** |
| *Crithidia/ Lotmaria* | A | 49 | 39 | 79.6 | 66.3 | 89.5 | 5.39 | 6.77 | 8.84 |
| | M | 28 | 25 | 89.3 | 72.2 | 97 | 6.02 | 6.75 | 8.48 |
| | F | 21 | 14 | 66.7 | 44.9 | 84.8 | 4.55 | 6.82 | 8.84 |
| *Nosema apis* | A | 49 | 6 | 12.2 | 5.5 | 24.1 | 0.59 | 4.82 | 9.37 |
| | M | 28 | 3 | 10.7 | 3 | 27.8 | 0.34 | 3.19 | 3.49 |
| | F | 21 | 3 | 14.3 | 4 | 35.1 | 0.92 | 6.45 | 9.37 |
| *Nosema ceranae* | A | 49 | 46 | 93.9 | 83.3 | 98.3 | 6.31 | 6.73 | 9.54 |
| | M | 28 | 26 | 92.9 | 77.6 | 98.7 | 6.4 | 6.89 | 9.54 |
| | F | 21 | 20 | 95.2 | 77.3 | 99.8 | 6.2 | 6.51 | 9.45 |
| **Bacteria** | | | | | | | | | |
| *Melissococcus plutonius* | A | 49 | 2 | 4.1 | 0.7 | 13.6 | 0.14 | 3.33 | 3.43 |
| | M | 28 | 1 | 3.6 | 0.2 | 17 | 0.12 | 3.43 | 3.43 |
| | F | 21 | 1 | 4.8 | 0.2 | 22.7 | 0.15 | 3.22 | 3.22 |
| *Paenibacillus larvae* | A | 49 | 1 | 2 | 0.1 | 10.5 | 0.08 | 3.68 | 3.68 |
| | M | 28 | 1 | 3.6 | 0.2 | 17 | 0.13 | 3.68 | 3.68 |
| | F | 21 | 0 | 0 | 0 | 15.2 | 0 | NA | 0 |
| **Viruses** | | | | | | | | | |
| Acute bee paralysis virus | A | 49 | 38 | 77.6 | 63.5 | 87.4 | 4.61 | 5.94 | 7.9 |
| | M | 28 | 20 | 71.4 | 51.8 | 85.8 | 3.96 | 5.54 | 7.61 |
| | F | 21 | 18 | 85.7 | 64.9 | 96 | 5.47 | 6.39 | 7.9 |
| Black queen cell virus | A | 49 | 44 | 89.8 | 78.1 | 95.9 | 4.9 | 5.46 | 7.71 |
| | M | 28 | 25 | 89.3 | 72.2 | 97 | 4.89 | 5.48 | 7.71 |
| | F | 21 | 19 | 90.5 | 69.9 | 98.3 | 4.92 | 5.44 | 7.52 |
| Chronic bee paralysis virus | A | 49 | 3 | 6.1 | 1.7 | 16.7 | 0.31 | 5.06 | 7.9 |
| | M | 28 | 3 | 10.7 | 3 | 27.8 | 0.54 | 5.06 | 7.9 |
| | F | 21 | 0 | 0 | 0 | 15.2 | 0 | NA | 0 |
| Deformed wing virus A | A | 49 | 5 | 10.2 | 4.1 | 21.9 | 0.45 | 4.38 | 4.57 |
| | M | 28 | 4 | 14.3 | 5 | 31.6 | 0.63 | 4.41 | 4.57 |
| | F | 21 | 1 | 4.8 | 0.2 | 22.7 | 0.2 | 4.27 | 4.27 |
| Deformed wing virus B | A | 49 | 10 | 20.4 | 10.5 | 33.7 | 1.41 | 6.92 | 8.42 |
| | M | 28 | 7 | 25 | 11.4 | 44.5 | 1.67 | 6.67 | 8.42 |
| | F | 21 | 3 | 14.3 | 4 | 35.1 | 1.07 | 7.5 | 7.91 |
| Israeli acute paralysis virus | A | 49 | 2 | 4.1 | 0.7 | 13.6 | 0.05 | 1.34 | 1.45 |
| | M | 28 | 0 | 0 | 0 | 11.4 | 0 | NA | 0 |
| | F | 21 | 2 | 9.5 | 1.7 | 30.1 | 0.13 | 1.34 | 1.45 |
| Lake Sinai virus | A | 49 | 46 | 93.9 | 83.3 | 98.3 | 5.24 | 5.59 | 7.58 |
| | M | 28 | 28 | 100 | 88.6 | 100 | 5.73 | 5.73 | 7.58 |
| | F | 21 | 18 | 85.7 | 64.9 | 96 | 4.59 | 5.36 | 6.98 |
| Sacbrood virus | A | 49 | 27 | 55.1 | 40.6 | 68.7 | 2.3 | 4.18 | 6.06 |
| | M | 28 | 18 | 64.3 | 44.5 | 80.8 | 2.73 | 4.24 | 6.06 |
| | F | 21 | 9 | 42.9 | 22.7 | 64.9 | 1.73 | 4.04 | 5.57 |
| Slow bee paralysis virus | A | 49 | 3 | 6.1 | 1.7 | 16.7 | 0.18 | 2.92 | 4.49 |
| | M | 28 | 2 | 7.1 | 1.3 | 22.4 | 0.25 | 3.49 | 4.49 |
| | F | 21 | 1 | 4.8 | 0.2 | 22.7 | 0.08 | 1.78 | 1.78 |
| Varroa destructor macula-like virus | A | 49 | 1 | 2 | 0.1 | 10.5 | 0.05 | 2.28 | 2.28 |
| | M | 28 | 1 | 3.6 | 0.2 | 17 | 0.08 | 2.28 | 2.28 |
| | F | 21 | 0 | 0 | 0 | 15.2 | 0 | NA | 0 |

**Table S8:** Overview of prevalence and colony-level loads of 15 microparasites for study region 2 (Coburg & Lichtenfels). Three other parasites assayed in this study (*Acarapis woodi*, Invertebrate iridescent virus 6, and Kashmir bee virus) were not detected in any of the four study regions. A: all colonies, M: managed colonies, and F: feral colonies.

| | | | | **Prevalence (%)** | | | **$Log_{10}$ (n / 100 ng RNA+1)** | | |
|---|---|---|---|---|---|---|---|---|---|
| **Parasite Taxon** | **Sample** | **N** | **N (positives)** | **Estimate** | **Lower CI** | **Upper CI** | **Mean** | **Mean (positives)** | **Max** |
| *Crithidia/ Lotmaria* | A | 33 | 28 | 84.8 | 68.7 | 93.8 | 5.92 | 6.98 | 8.59 |
| | M | 17 | 15 | 88.2 | 65.4 | 97.9 | 6.24 | 7.07 | 8.59 |
| | F | 16 | 13 | 81.3 | 56.6 | 94.7 | 5.59 | 6.88 | 8.51 |
| *Nosema apis* | A | 33 | 0 | 0 | 0 | 9.6 | 0 | NA | 0 |
| | M | 17 | 0 | 0 | 0 | 18.9 | 0 | NA | 0 |
| | F | 16 | 0 | 0 | 0 | 20.1 | 0 | NA | 0 |
| *Nosema ceranae* | A | 33 | 33 | 100 | 90.4 | 100 | 8.28 | 8.28 | 9.59 |
| | M | 17 | 17 | 100 | 81.1 | 100 | 8.65 | 8.65 | 9.51 |
| | F | 16 | 16 | 100 | 79.9 | 100 | 7.88 | 7.88 | 9.59 |
| **Bacteria** | | | | | | | | | |
| *Melissococcus plutonius* | A | 33 | 3 | 9.1 | 2.5 | 23.6 | 0.21 | 2.27 | 2.48 |
| | M | 17 | 0 | 0 | 0 | 18.9 | 0 | NA | 0 |
| | F | 16 | 3 | 18.8 | 5.3 | 43.4 | 0.43 | 2.27 | 2.48 |
| *Paenibacillus larvae* | A | 33 | 0 | 0 | 0 | 9.6 | 0 | NA | 0 |
| | M | 17 | 0 | 0 | 0 | 18.9 | 0 | NA | 0 |
| | F | 16 | 0 | 0 | 0 | 20.1 | 0 | NA | 0 |
| **Viruses** | | | | | | | | | |
| Acute bee paralysis virus | A | 33 | 2 | 6.1 | 1.1 | 19.2 | 0.15 | 2.47 | 2.69 |
| | M | 17 | 2 | 11.8 | 2.1 | 34.6 | 0.29 | 2.47 | 2.69 |
| | F | 16 | 0 | 0 | 0 | 20.1 | 0 | NA | 0 |
| Black queen cell virus | A | 33 | 33 | 100 | 90.4 | 100 | 5.35 | 5.35 | 8.13 |
| | M | 17 | 17 | 100 | 81.1 | 100 | 5.28 | 5.28 | 8.13 |
| | F | 16 | 16 | 100 | 79.9 | 100 | 5.43 | 5.43 | 7.31 |
| Chronic bee paralysis virus | A | 33 | 4 | 12.1 | 4.2 | 28.2 | 0.75 | 6.21 | 7.49 |
| | M | 17 | 4 | 23.5 | 8.5 | 48.9 | 1.46 | 6.21 | 7.49 |
| | F | 16 | 0 | 0 | 0 | 20.1 | 0 | NA | 0 |
| Deformed wing virus A | A | 33 | 5 | 15.2 | 6.2 | 31.3 | 0.7 | 4.61 | 4.9 |
| | M | 17 | 5 | 29.4 | 12.4 | 54.4 | 1.36 | 4.61 | 4.9 |
| | F | 16 | 0 | 0 | 0 | 20.1 | 0 | NA | 0 |
| Deformed wing virus B | A | 33 | 10 | 30.3 | 15.7 | 48.5 | 1.48 | 4.89 | 7 |
| | M | 17 | 6 | 35.3 | 16.4 | 59.4 | 1.69 | 4.8 | 7 |
| | F | 16 | 4 | 25 | 9 | 50 | 1.26 | 5.03 | 5.57 |
| Israeli acute paralysis virus | A | 33 | 0 | 0 | 0 | 9.6 | 0 | NA | 0 |
| | M | 17 | 0 | 0 | 0 | 18.9 | 0 | NA | 0 |
| | F | 16 | 0 | 0 | 0 | 20.1 | 0 | NA | 0 |
| Lake Sinai virus | A | 33 | 31 | 93.9 | 80.8 | 98.9 | 5.55 | 5.91 | 7.34 |
| | M | 17 | 15 | 88.2 | 65.4 | 97.9 | 5.1 | 5.78 | 7.18 |
| | F | 16 | 16 | 100 | 79.9 | 100 | 6.03 | 6.03 | 7.34 |
| Sacbrood virus | A | 33 | 11 | 33.3 | 19 | 51.6 | 1.66 | 4.98 | 6.98 |
| | M | 17 | 4 | 23.5 | 8.5 | 48.9 | 1.1 | 4.68 | 5.83 |
| | F | 16 | 7 | 43.8 | 20.1 | 70 | 2.25 | 5.15 | 6.98 |
| Slow bee paralysis virus | A | 33 | 0 | 0 | 0 | 9.6 | 0 | NA | 0 |
| | M | 17 | 0 | 0 | 0 | 18.9 | 0 | NA | 0 |
| | F | 16 | 0 | 0 | 0 | 20.1 | 0 | NA | 0 |
| Varroa destructor macula-like virus | A | 33 | 0 | 0 | 0 | 9.6 | 0 | NA | 0 |
| | M | 17 | 0 | 0 | 0 | 18.9 | 0 | NA | 0 |
| | F | 16 | 0 | 0 | 0 | 20.1 | 0 | NA | 0 |

**Table S9:** Overview of prevalence and colony-level loads of 15 microparasites for study region 3 (Weilheim-Schongau). Three other parasites assayed in this study (*Acarapis woodi*, Invertebrate iridescent virus 6, and Kashmir bee virus) were not detected in any of the four study regions. A: all colonies, M: managed colonies, and F: feral colonies.

| | | | | Prevalence (%) | | | $Log_{10}$ (n / 100 ng RNA+1) | | |
|---|---|---|---|---|---|---|---|---|---|
| **Parasite Taxon** | **Sample** | **N** | **N (positives)** | **Estimate** | **Lower CI** | **Upper CI** | **Mean** | **Mean (positives)** | **Max** |
| *Crithidia/ Lotmaria* | A | 24 | 24 | 100 | 86.7 | 100 | 7.73 | 7.73 | 8.75 |
| | M | 12 | 12 | 100 | 76.4 | 100 | 7.83 | 7.83 | 8.75 |
| | F | 12 | 12 | 100 | 76.4 | 100 | 7.62 | 7.62 | 8.55 |
| *Nosema apis* | A | 24 | 0 | 0 | 0 | 13.3 | 0 | NA | 0 |
| | M | 12 | 0 | 0 | 0 | 23.6 | 0 | NA | 0 |
| | F | 12 | 0 | 0 | 0 | 23.6 | 0 | NA | 0 |
| *Nosema ceranae* | A | 24 | 22 | 91.7 | 73.8 | 98.5 | 5.25 | 5.73 | 9.35 |
| | M | 12 | 11 | 91.7 | 63.4 | 99.6 | 5.47 | 5.96 | 9.35 |
| | F | 12 | 11 | 91.7 | 63.4 | 99.6 | 5.04 | 5.49 | 8.85 |
| **Bacteria** | | | | | | | | | |
| *Melissococcus plutonius* | A | 24 | 5 | 20.8 | 8.6 | 41.4 | 0.56 | 2.68 | 3.71 |
| | M | 12 | 4 | 33.3 | 12.3 | 63.4 | 0.99 | 2.96 | 3.71 |
| | F | 12 | 1 | 8.3 | 0.4 | 36.6 | 0.13 | 1.58 | 1.58 |
| *Paenibacillus larvae* | A | 24 | 0 | 0 | 0 | 13.3 | 0 | NA | 0 |
| | M | 12 | 0 | 0 | 0 | 23.6 | 0 | NA | 0 |
| | F | 12 | 0 | 0 | 0 | 23.6 | 0 | NA | 0 |
| **Viruses** | | | | | | | | | |
| Acute bee paralysis virus | A | 24 | 14 | 58.3 | 37 | 76.6 | 2.47 | 4.24 | 8.12 |
| | M | 12 | 9 | 75 | 45.6 | 92.8 | 3.95 | 5.26 | 8.12 |
| | F | 12 | 5 | 41.7 | 18.1 | 70.6 | 1 | 2.41 | 2.8 |
| Black queen cell virus | A | 24 | 23 | 95.8 | 80.2 | 99.8 | 5.84 | 6.09 | 8.55 |
| | M | 12 | 12 | 100 | 76.4 | 100 | 6.27 | 6.27 | 8.55 |
| | F | 12 | 11 | 91.7 | 63.4 | 99.6 | 5.4 | 5.89 | 8.3 |
| Chronic bee paralysis virus | A | 24 | 0 | 0 | 0 | 13.3 | 0 | NA | 0 |
| | M | 12 | 0 | 0 | 0 | 23.6 | 0 | NA | 0 |
| | F | 12 | 0 | 0 | 0 | 23.6 | 0 | NA | 0 |
| Deformed wing virus A | A | 24 | 0 | 0 | 0 | 13.3 | 0 | NA | 0 |
| | M | 12 | 0 | 0 | 0 | 23.6 | 0 | NA | 0 |
| | F | 12 | 0 | 0 | 0 | 23.6 | 0 | NA | 0 |
| Deformed wing virus B | A | 24 | 14 | 58.3 | 37 | 76.6 | 3.6 | 6.17 | 8.45 |
| | M | 12 | 8 | 66.7 | 36.6 | 87.7 | 4.39 | 6.58 | 8.45 |
| | F | 12 | 6 | 50 | 23.4 | 76.6 | 2.81 | 5.62 | 8.41 |
| Israeli acute paralysis virus | A | 24 | 0 | 0 | 0 | 13.3 | 0 | NA | 0 |
| | M | 12 | 0 | 0 | 0 | 23.6 | 0 | NA | 0 |
| | F | 12 | 0 | 0 | 0 | 23.6 | 0 | NA | 0 |
| Lake Sinai virus | A | 24 | 18 | 75 | 54.3 | 88.5 | 3.59 | 4.79 | 6.79 |
| | M | 12 | 9 | 75 | 45.6 | 92.8 | 3.34 | 4.45 | 6.77 |
| | F | 12 | 9 | 75 | 45.6 | 92.8 | 3.84 | 5.12 | 6.79 |
| Sacbrood virus | A | 24 | 13 | 54.2 | 33.9 | 73.8 | 2.44 | 4.5 | 6.11 |
| | M | 12 | 8 | 66.7 | 36.6 | 87.7 | 3.11 | 4.67 | 5.59 |
| | F | 12 | 5 | 41.7 | 18.1 | 70.6 | 1.76 | 4.22 | 6.11 |
| Slow bee paralysis virus | A | 24 | 0 | 0 | 0 | 13.3 | 0 | NA | 0 |
| | M | 12 | 0 | 0 | 0 | 23.6 | 0 | NA | 0 |
| | F | 12 | 0 | 0 | 0 | 23.6 | 0 | NA | 0 |
| Varroa destructor macula-like virus | A | 24 | 0 | 0 | 0 | 13.3 | 0 | NA | 0 |
| | M | 12 | 0 | 0 | 0 | 23.6 | 0 | NA | 0 |
| | F | 12 | 0 | 0 | 0 | 23.6 | 0 | NA | 0 |

**Table S10:** Overview of prevalence and colony-level loads of 15 microparasites for study region 4 (Munich). Three other parasites assayed in this study (*Acarapis woodi*, Invertebrate iridescent virus 6, and Kashmir bee virus) were not detected in any of the four study regions. A: all colonies, M: managed colonies, and F: feral colonies.

| | | | | **Prevalence (%)** | | | **$Log_{10}$ (n / 100 ng RNA+1)** | | |
|---|---|---|---|---|---|---|---|---|---|
| **Parasite Taxon** | **Sample** | **N** | **N (positives)** | **Estimate** | **Lower CI** | **Upper CI** | **Mean** | **Mean (positives)** | **Max** |
| *Crithidia/ Lotmaria* | A | 32 | 26 | 81.3 | 64.4 | 91.5 | 5.66 | 6.97 | 8.49 |
| | M | 17 | 15 | 88.2 | 65.4 | 97.9 | 6.06 | 6.86 | 8.49 |
| | F | 15 | 11 | 73.3 | 46.5 | 90.3 | 5.22 | 7.12 | 8.14 |
| *Nosema apis* | A | 32 | 0 | 0 | 0 | 9.9 | 0 | NA | 0 |
| | M | 17 | 0 | 0 | 0 | 18.9 | 0 | NA | 0 |
| | F | 15 | 0 | 0 | 0 | 21.5 | 0 | NA | 0 |
| *Nosema ceranae* | A | 32 | 32 | 100 | 90.1 | 100 | 6.12 | 6.12 | 9.57 |
| | M | 17 | 17 | 100 | 81.1 | 100 | 6.12 | 6.12 | 9.57 |
| | F | 15 | 15 | 100 | 78.5 | 100 | 6.13 | 6.13 | 9.35 |
| **Bacteria** | | | | | | | | | |
| *Melissococcus plutonius* | A | 32 | 1 | 3.1 | 0.2 | 16.2 | 0.1 | 3.34 | 3.34 |
| | M | 17 | 1 | 5.9 | 0.3 | 28.2 | 0.2 | 3.34 | 3.34 |
| | F | 15 | 0 | 0 | 0 | 21.5 | 0 | NA | 0 |
| *Paenibacillus larvae* | A | 32 | 0 | 0 | 0 | 9.9 | 0 | NA | 0 |
| | M | 17 | 0 | 0 | 0 | 18.9 | 0 | NA | 0 |
| | F | 15 | 0 | 0 | 0 | 21.5 | 0 | NA | 0 |
| **Viruses** | | | | | | | | | |
| Acute bee paralysis virus | A | 32 | 15 | 46.9 | 29.5 | 64.4 | 2.61 | 5.57 | 7.81 |
| | M | 17 | 9 | 52.9 | 28.2 | 74.7 | 3.32 | 6.28 | 7.81 |
| | F | 15 | 6 | 40 | 18.6 | 66.8 | 1.8 | 4.5 | 7.64 |
| Black queen cell virus | A | 32 | 31 | 96.9 | 83.8 | 99.8 | 6.11 | 6.3 | 7.93 |
| | M | 17 | 16 | 94.1 | 71.8 | 99.7 | 6.23 | 6.62 | 7.88 |
| | F | 15 | 15 | 100 | 78.5 | 100 | 5.97 | 5.97 | 7.93 |
| Chronic bee paralysis virus | A | 32 | 5 | 15.6 | 6.4 | 32.3 | 1.03 | 6.6 | 7.93 |
| | M | 17 | 4 | 23.5 | 8.5 | 48.9 | 1.49 | 6.32 | 7.93 |
| | F | 15 | 1 | 6.7 | 0.3 | 30.2 | 0.51 | 7.71 | 7.71 |
| Deformed wing virus A | A | 32 | 6 | 18.8 | 8.5 | 35.6 | 1.09 | 5.82 | 9.28 |
| | M | 17 | 5 | 29.4 | 12.4 | 54.4 | 1.79 | 6.1 | 9.28 |
| | F | 15 | 1 | 6.7 | 0.3 | 30.2 | 0.3 | 4.45 | 4.45 |
| Deformed wing virus B | A | 32 | 24 | 75 | 57.7 | 87.8 | 4.97 | 6.62 | 8.63 |
| | M | 17 | 14 | 82.4 | 58.3 | 95 | 5.57 | 6.76 | 8.63 |
| | F | 15 | 10 | 66.7 | 39.4 | 85.8 | 4.29 | 6.43 | 8.27 |
| Israeli acute paralysis virus | A | 32 | 0 | 0 | 0 | 9.9 | 0 | NA | 0 |
| | M | 17 | 0 | 0 | 0 | 18.9 | 0 | NA | 0 |
| | F | 15 | 0 | 0 | 0 | 21.5 | 0 | NA | 0 |
| Lake Sinai virus | A | 32 | 32 | 100 | 90.1 | 100 | 6.46 | 6.46 | 7.48 |
| | M | 17 | 17 | 100 | 81.1 | 100 | 6.55 | 6.55 | 7.48 |
| | F | 15 | 15 | 100 | 78.5 | 100 | 6.36 | 6.36 | 7.41 |
| Sacbrood virus | A | 32 | 21 | 65.6 | 47.3 | 80.4 | 2.98 | 4.55 | 8.36 |
| | M | 17 | 10 | 58.8 | 34.6 | 81.1 | 2.72 | 4.63 | 8.36 |
| | F | 15 | 11 | 73.3 | 46.5 | 90.3 | 3.28 | 4.47 | 5.95 |
| Slow bee paralysis virus | A | 32 | 0 | 0 | 0 | 9.9 | 0 | NA | 0 |
| | M | 17 | 0 | 0 | 0 | 18.9 | 0 | NA | 0 |
| | F | 15 | 0 | 0 | 0 | 21.5 | 0 | NA | 0 |
| Varroa destructor macula-like virus | A | 32 | 4 | 12.5 | 4.4 | 28.2 | 0.33 | 2.66 | 3.02 |
| | M | 17 | 2 | 11.8 | 2.1 | 34.6 | 0.33 | 2.81 | 3.02 |
| | F | 15 | 2 | 13.3 | 2.4 | 39.4 | 0.33 | 2.51 | 2.91 |

**Table S11:** Result of a likelihood ratio test (function “anova()” in R) comparing two nested models explaining the number of detected parasites. Adding the factor “colony type” to the factor “region” significantly improves model fit.

| Model | Df | AIC | BIC | logLik | deviance | Chisq | Δ Df | Pr(>Chisq) |
|---|---|---|---|---|---|---|---|---|
| glmmTMB(Number of parasites ~ **region**, dispformula = ~region, family = genpois() | 8 | 472.38 | 495.798 | -228.190 | 456.38 | | | |
| glmmTMB(Number of parasites ~ **colony type + region**, dispformula = ~region, family = genpois() | 12 | 457.15 | 492.277 | -216.575 | 433.15 | 23.23 | 4 | 0.0001 |

**Table S12:** Result of a likelihood ratio test (function “anova()” in R) comparing two nested models of the number of detected parasites. The interaction between “colony type” and “region” does not significantly improve model fit.

| Model | Df | AIC | BIC | logLik | deviance | Chisq | Δ Df | Pr(>Chisq) |
|---|---|---|---|---|---|---|---|---|
| glmmTMB(Number of parasites ~ **colony type + region**, dispformula = ~region, family = genpois()) | 12 | 457.150 | 492.277 | -216.575 | 433.150 | | | |
| glmmTMB(Number of parasites ~ **colony type + region + colony type:region**, dispformula = ~region, family = genpois()) | 24 | 471.604 | 541.858 | -211.802 | 423.604 | 9.546 | 12 | 0.656 |

**Table S13:** Mean numbers of detected parasites per colony and the respective 95% confidence limits (CI) estimated by a generalized linear model [model formula: glmmTMB(Number of parasites ~ colony type + region + colony type:region, dispformula= ~ region, family = genpois()] for the five colony types and divided by study region.

| Region | Colony type | Estimate | Lower CI | Upper CI |
|---|---|---|---|---|
| Swabian Alb | Managed overwintered | 6.052 | 5.373 | 6.817 |
| | Managed hived swarm | 7.273 | 5.412 | 9.773 |
| | Managed nucleus | 5.587 | 4.875 | 6.403 |
| | Feral overwintered | 5.875 | 4.672 | 7.388 |
| | Feral founder | 5.423 | 4.836 | 6.080 |
| Coburg & Lichtenfels | Managed overwintered | 5.576 | 5.092 | 6.106 |
| | Managed hived swarm | 5.978 | 5.126 | 6.972 |
| | Managed nucleus | 4.280 | 3.718 | 4.928 |
| | Feral overwintered | 4.577 | 3.916 | 5.350 |
| | Feral founder | 4.681 | 4.290 | 5.108 |
| Weilheim-Schongau | Managed overwintered | 6.907 | 6.052 | 7.883 |
| | Managed hived swarm | 6.931 | 5.526 | 8.692 |
| | Managed nucleus | 6.529 | 5.516 | 7.728 |
| | Feral overwintered | 6.435 | 5.618 | 7.372 |
| | Feral founder | 5.328 | 4.603 | 6.168 |
| Munich | Managed overwintered | 6.956 | 6.253 | 7.738 |
| | Managed hived swarm | 6.461 | 5.177 | 8.065 |
| | Managed nucleus | 6.503 | 5.682 | 7.443 |
| | Feral overwintered | 5.857 | 4.912 | 6.983 |
| | Feral founder | 5.639 | 5.063 | 6.280 |

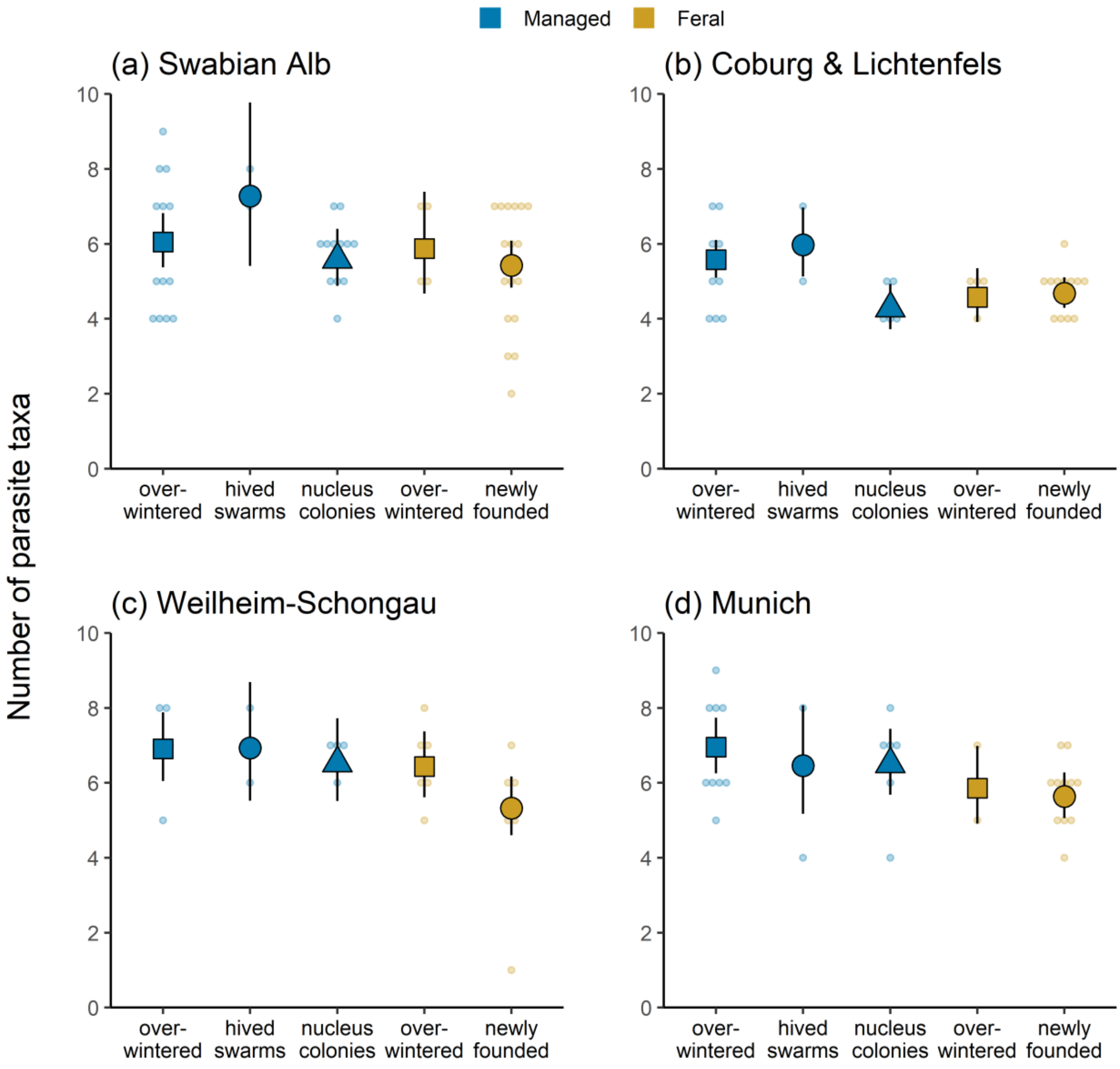


**Figure S2 (a)–(d):** Number of microparasite taxa detected among the 18 taxa assayed in relation to colony type for each of the four study regions. Dots are raw data; large symbols and vertical lines give model-estimated means and 95%-confidence intervals.

**Table S14:** Overview of prevalence and colony-level loads of 15 microparasites for five colony types. Three other parasites assayed in this study (*Acarapis woodi*, Invertebrate iridescent virus 6, and Kashmir bee virus) were not detected in any colony. MO: overwintered managed colonies, MS: managed hived swarms, MN: managed nucleus colonies, FO: overwintered feral colonies and FF: newly founded feral colonies.

| | | | | Prevalence (%) | | | Log10 (n / 100 ng RNA+1) | | |
|---|---|---|---|---|---|---|---|---|---|
| **Parasite Taxon** | **Colony type** | **N** | **N (positives)** | **Estimate** | **Lower CI** | **Upper CI** | **Mean** | **Mean (positives)** | **Max** |
| *Crithidia/ Lotmaria* | MO | 38 | 36 | 94.7 | 82.4 | 99.1 | 6.92 | 7.31 | 8.75 |
| | MS | 9 | 7 | 77.8 | 44 | 95.9 | 6.15 | 7.91 | 8.59 |
| | MN | 27 | 24 | 88.9 | 71.1 | 96.9 | 5.68 | 6.39 | 8.39 |
| | FO | 18 | 14 | 77.8 | 52.9 | 92 | 5.92 | 7.61 | 8.35 |
| | FF | 46 | 36 | 78.3 | 64.4 | 88.8 | 5.4 | 6.89 | 8.84 |
| *Nosema apis* | MO | 38 | 2 | 5.3 | 0.9 | 17.6 | 0.16 | 3.04 | 3.2 |
| | MS | 9 | 1 | 11.1 | 0.6 | 44.3 | 0.39 | 3.49 | 3.49 |
| | MN | 27 | 0 | 0 | 0 | 11.8 | 0 | NA | 0 |
| | FO | 18 | 1 | 5.6 | 0.3 | 26.6 | 0.16 | 2.94 | 2.94 |
| | FF | 46 | 2 | 4.3 | 0.8 | 14.5 | 0.36 | 8.21 | 9.37 |
| *Nosema ceranae* | MO | 38 | 37 | 97.4 | 86.4 | 99.9 | 7.06 | 7.26 | 9.57 |
| | MS | 9 | 9 | 100 | 68.4 | 100 | 7.51 | 7.51 | 9.54 |
| | MN | 27 | 25 | 92.6 | 76.7 | 98.7 | 5.92 | 6.39 | 9.51 |
| | FO | 18 | 18 | 100 | 82.2 | 100 | 6.58 | 6.58 | 9.51 |
| | FF | 46 | 44 | 95.7 | 85.5 | 99.2 | 6.31 | 6.6 | 9.59 |
| **Bacteria** | | | | | | | | | |
| *Melissococcus plutonius* | MO | 38 | 4 | 10.5 | 3.7 | 24.4 | 0.35 | 3.33 | 3.71 |
| | MS | 9 | 1 | 11.1 | 0.6 | 44.3 | 0.26 | 2.3 | 2.3 |
| | MN | 27 | 1 | 3.7 | 0.2 | 17.6 | 0.11 | 2.96 | 2.96 |
| | FO | 18 | 1 | 5.6 | 0.3 | 26.6 | 0.09 | 1.58 | 1.58 |
| | FF | 46 | 4 | 8.7 | 3 | 20.1 | 0.22 | 2.51 | 3.22 |
| *Paenibacillus larvae* | MO | 38 | 1 | 2.6 | 0.1 | 13.6 | 0.1 | 3.68 | 3.68 |
| | MS | 9 | 0 | 0 | 0 | 31.6 | 0 | NA | 0 |
| | MN | 27 | 0 | 0 | 0 | 11.8 | 0 | NA | 0 |
| | FO | 18 | 0 | 0 | 0 | 17.8 | 0 | NA | 0 |
| | FF | 46 | 0 | 0 | 0 | 6.9 | 0 | NA | 0 |
| **Viruses** | | | | | | | | | |
| Acute bee paralysis virus | MO | 38 | 19 | 50 | 33.8 | 66.2 | 2.83 | 5.66 | 7.61 |
| | MS | 9 | 6 | 66.7 | 31.6 | 90.2 | 3.52 | 5.29 | 8.12 |
| | MN | 27 | 15 | 55.6 | 36.6 | 73.1 | 2.98 | 5.36 | 7.81 |
| | FO | 18 | 9 | 50 | 26.6 | 73.4 | 2.52 | 5.03 | 7.9 |
| | FF | 46 | 20 | 43.5 | 29.2 | 58.9 | 2.36 | 5.44 | 7.87 |
| Black queen cell virus | MO | 38 | 34 | 89.5 | 75.6 | 96.3 | 5.18 | 5.79 | 7.88 |
| | MS | 9 | 9 | 100 | 68.4 | 100 | 5.56 | 5.56 | 8.05 |
| | MN | 27 | 27 | 100 | 88.2 | 100 | 5.96 | 5.96 | 8.55 |
| | FO | 18 | 18 | 100 | 82.2 | 100 | 5.84 | 5.84 | 8.3 |
| | FF | 46 | 43 | 93.5 | 82.2 | 98.2 | 5.21 | 5.57 | 7.93 |
| Chronic bee paralysis virus | MO | 38 | 7 | 18.4 | 8.4 | 33.8 | 1.26 | 6.86 | 7.93 |
| | MS | 9 | 3 | 33.3 | 9.8 | 68.4 | 1.4 | 4.2 | 7.33 |
| | MN | 27 | 1 | 3.7 | 0.2 | 17.6 | 0.17 | 4.66 | 4.66 |
| | FO | 18 | 1 | 5.6 | 0.3 | 26.6 | 0.43 | 7.71 | 7.71 |
| | FF | 46 | 0 | 0 | 0 | 6.9 | 0 | NA | 0 |
| Deformed wing virus A | MO | 38 | 8 | 21.1 | 10 | 36.5 | 1.15 | 5.45 | 9.28 |
| | MS | 9 | 1 | 11.1 | 0.6 | 44.3 | 0.54 | 4.9 | 4.9 |
| | MN | 27 | 5 | 18.5 | 7.6 | 36.7 | 0.84 | 4.53 | 4.65 |
| | FO | 18 | 0 | 0 | 0 | 17.8 | 0 | NA | 0 |
| | FF | 46 | 2 | 4.3 | 0.8 | 14.5 | 0.19 | 4.36 | 4.45 |

**Table S14** (*continued*).

| Parasite Taxon | Sample | N | N (positives) | Prevalence (%) Estimate | Lower CI | Upper CI | Log10 (n / 100 ng RNA+1) Mean | Mean (positives) | Max |
|---|---|---|---|---|---|---|---|---|---|
| Deformed wing virus B | MO | 38 | 20 | 52.6 | 36.5 | 68.9 | 3.32 | 6.31 | 8.63 |
| | MS | 9 | 5 | 55.6 | 25.1 | 83.1 | 3.5 | 6.3 | 7.96 |
| | MN | 27 | 10 | 37 | 20.2 | 57.1 | 2.41 | 6.52 | 8.45 |
| | FO | 18 | 11 | 61.1 | 37.5 | 82.2 | 3.68 | 6.02 | 7.91 |
| | FF | 46 | 12 | 26.1 | 14.5 | 41.1 | 1.62 | 6.2 | 8.41 |
| Israeli acute paralysis virus | MO | 38 | 0 | 0 | 0 | 8.4 | 0 | NA | 0 |
| | MS | 9 | 0 | 0 | 0 | 31.6 | 0 | NA | 0 |
| | MN | 27 | 0 | 0 | 0 | 11.8 | 0 | NA | 0 |
| | FO | 18 | 0 | 0 | 0 | 17.8 | 0 | NA | 0 |
| | FF | 46 | 2 | 4.3 | 0.8 | 14.5 | 0.06 | 1.34 | 1.45 |
| Lake Sinai virus | MO | 38 | 36 | 94.7 | 82.4 | 99.1 | 5.34 | 5.64 | 7.58 |
| | MS | 9 | 8 | 88.9 | 55.7 | 99.4 | 5.52 | 6.21 | 7.07 |
| | MN | 27 | 25 | 92.6 | 76.7 | 98.7 | 5.4 | 5.83 | 7.39 |
| | FO | 18 | 17 | 94.4 | 73.4 | 99.7 | 5.95 | 6.3 | 7.31 |
| | FF | 46 | 41 | 89.1 | 76.6 | 95.6 | 4.94 | 5.54 | 7.41 |
| Sacbrood virus | MO | 38 | 19 | 50 | 33.8 | 66.2 | 2.19 | 4.38 | 6.99 |
| | MS | 9 | 4 | 44.4 | 16.9 | 74.9 | 1.73 | 3.88 | 4.48 |
| | MN | 27 | 17 | 63 | 42.9 | 79.8 | 2.96 | 4.71 | 8.36 |
| | FO | 18 | 8 | 44.4 | 23.6 | 67.4 | 2 | 4.5 | 6.11 |
| | FF | 46 | 24 | 52.2 | 37.8 | 66.6 | 2.32 | 4.44 | 6.98 |
| Slow bee paralysis virus | MO | 38 | 2 | 5.3 | 0.9 | 17.6 | 0.18 | 3.49 | 4.49 |
| | MS | 9 | 0 | 0 | 0 | 31.6 | 0 | NA | 0 |
| | MN | 27 | 0 | 0 | 0 | 11.8 | 0 | NA | 0 |
| | FO | 18 | 0 | 0 | 0 | 17.8 | 0 | NA | 0 |
| | FF | 46 | 1 | 2.2 | 0.1 | 11.2 | 0.04 | 1.78 | 1.78 |
| Varroa destructor macula-like virus | MO | 38 | 2 | 5.3 | 0.9 | 17.6 | 0.15 | 2.81 | 3.02 |
| | MS | 9 | 0 | 0 | 0 | 31.6 | 0 | NA | 0 |
| | MN | 27 | 1 | 3.7 | 0.2 | 17.6 | 0.08 | 2.28 | 2.28 |
| | FO | 18 | 1 | 5.6 | 0.3 | 26.6 | 0.12 | 2.11 | 2.11 |
| | FF | 46 | 1 | 2.2 | 0.1 | 11.2 | 0.06 | 2.91 | 2.91 |
